# Antiphospholipid syndrome (APS) is a platelet factor 4 (PF4)-centric immunothrombotic disorder

**DOI:** 10.1101/2025.11.07.687095

**Authors:** Conroy O. Field, Amrita Sarkar, Khalil Bdeir, Hyunjun Kim, Santosh K. Yadav, Jenna Oberg, Guohua Zhou, Steven Jiang, Manas Gumedelli, Keith R. McCrae, Gowthami M. Arepally, Jerome Rollin, Yves Gruel, Thomas L. Ortel, M. Anna Kowalska, Douglas B. Cines, Kandace Gollomp, Lubica Rauova, Mortimer Poncz

## Abstract

Antiphospholipid syndrome (APS) is an immunothrombotic disorder, frequently attributed to autoantibodies that bind β2-glycoprotein I (β2GPI). A study showed that the platelet-specific chemokine, platelet factor 4 (PF4), binds to β2GPI, enhancing recognition of β2GPI by APS antibodies. APS antibodies induce the release of neutrophil extracellular traps (NETs), webs of decondensed chromatin that bind both PF4 and β2GPI. We propose that PF4 bridges β2GPI to NETs (and other PF4-targeted polyanions), leading to the formation of prothrombotic PF4:β2GPI:NET immunotargets in APS. Dynamic light-scattering studies of isolated IgGs from four patients with triple-positive APS show formation of PF4:β2GPI:NET complexes that bind APS antibodies. NETs released in a microfluidic system bound β2GPI, but only in the presence of PF4, forming a multimolecular APS antigenic target. Whole blood infused through a photochemically-injured, endothelium-lined microfluidic channel formed platelet-, fibrin-, and complement-rich thrombi that bound APS antibody only in the presence of PF4. Thrombi were reduced in size if either ADAMTS13 or DNase1 was infused. In a murine APS model, wildtype and transgenic mice expressing platelet human PF4 ± FcγRIIA developed more intense neutrophil rolling along veins, and more extensive thrombus formation following laser injury to cremaster arterioles and venules, whereas mice lacking PF4 did not. Three antigenically distinct anti-hPF4 monoclonal antibodies blocked thrombosis *in vitro*, and neutrophil rolling and thrombosis *in vivo*. Our studies provide new insights into the basis of APS that has mechanistic parallels to other known PF4 immunothrombotic disorders and offer potential diagnostic and non-anticoagulant therapeutic strategies for clinical management.

**Key points:**

- PF4 enhances β2GPI binding to NETs and these complexes are central to APS immunothrombosis.
- Anti-PF4 monoclonal antibodies block APS immunothrombosis in microfluidic and murine studies.

## Introduction

Antiphospholipid syndrome (APS) is a complex, acquired immunothrombotic disorder with a prevalence of ∼1:2,000 in the United States^1,2^. Patients may have macro- and/or micro-thrombotic events and obstetrical complications^3,4^. Diagnosis of APS requires having abnormalities in at least one of three laboratory tests: (*i*) a lupus anticoagulant, (*ii*) elevated IgG or IgM anti-cardiolipin antibodies, or (*iii*) elevated anti-β2-glycoprotein I (β2GPI) antibodies, confirmed on retesting at least 12 weeks apart^5^. The risk of thrombosis in APS is greatest in “triple-positive” patients^6,7^. β2GPI is a five-domain protein comprised of four similar complement control protein-like domains and a distinct domain V containing a large lysine-rich loop thought to be involved in binding to phospholipids^8^. The biological role of β2GPI is unclear, though it has been shown to influence complement and hemostatic function^9^. Studies of a β2GPI knockout mouse suggest that β2GPI may have anti-thrombotic effects^10^, but how that is linked mechanistically to the prothrombotic nature of APS has not been elucidated.

In a recent review of the potential biology of β2GPI, over 12 pathways were discussed^9^. Not mentioned was a 2010 paper^11^ describing β2GPI binding to platelet factor 4 (PF4, also known as CXCL4), a tetrameric chemokine abundantly expressed by megakaryocytes and stored in platelet α granules^12^. PF4 binding appeared to enhance the binding of APS antibodies to β2GPI, and the authors proposed – but did not test – that this duplex might be related to the prothrombotic nature of APS^11^. Recently, the anti-PF4 immunothrombotic disorder, heparin-induced thrombocytopenia (HIT), was recognized to be part of a larger family of immunothrombotic disorders with a similar mechanistic basis as vaccine-induced immune thrombotic thrombocytopenia (VITT) and the thrombotic complications occurring after natural adenoviral infection^13^. In these diseases, PF4 complexed to polyanions, including neutrophil extracellular traps (NETs) released in abundance from activated neutrophils at sites of thromboinflammation, serve as the immunothrombotic target and/or enhance thrombosis^14^. Tetrameric PF4 has an equatorial ring of positive charges. We noted previously that more than one polyanion can be cross-bound electrostatically by tetrameric PF4^15^. Video studies by our group showed that released NETs on their own appear fluffy, only becoming condensed into recognized NET strands upon exposure to and cross-binding by PF4^16^.

It has been shown that APS antibodies activate neutrophils, releasing NETs^17^, and that β2GPI bound to NETs may be important in APS thrombosis^18^. We posited that it is PF4:β2GPI:NET complexes that are central to thrombus development in APS. We confirmed by dynamic light scattering (DLS) that PF4 binds β2GPI, but also that higher molecular weight complexes were formed by PF4 plus β2GPI in the presence of DNA, the polyanion backbone of NETs, and that APS antibodies from triple-positive individuals bound to these PF4:β2GPI:DNA complexes. In a NET-lined microfluidic system, we found that binding of β2GPI to NETs, and the subsequent binding of anti-β2GPI antibodies, was low unless PF4 was included. In a photochemical-injured human umbilical vein endothelial cell (HUVEC)-lined system, we showed that PF4 needed to be present for vigorous platelet-, fibrin-, and complement-rich thrombi to form following the addition of APS antibodies to whole blood from healthy controls. In a passive antibody infusion murine model of APS, venous neutrophil rolling and adhesion, and venule and arteriole thrombosis in a cremaster laser injury system depended on the mice expressing either murine (m) or human (h) PF4. Moreover, three distinct monoclonal antibodies (moAbs), which blocked hPF4 biology differently, prevented β2GPI:hPF4:NET:APS immune complex formation in the DLS system. Anti-PF4 moAbs blocked thrombosis in the HUVEC microfluidic system and prevented slowing of venule neutrophil rolling and the prothrombotic state seen after inducing APS in murine models. The clinical implications of these findings, as well as their potential import for the physiologic role of β2GPI and PF4, are discussed.

## Materials and methods

### Human samples

Plasma from four anonymized patients with triple-positive APS and a history of thrombosis was studied (**Table 1**). IgG was isolated from plasma samples with Protein G agarose (ThermoFisher) as per the manufacturer’s instructions. Control plasma and IgGs were pooled plasma samples collected from ten healthy individuals without a history of thrombosis obtained at the Children’s Hospital of Philadelphia or from anonymized healthy volunteers at the University of Pennsylvania after approval by the appropriate institutional human subjects review board. Samples from patients with APS were studied with the approval of the institutional human subjects review board of Duke University Medical Center or Case Western Reserve University. All human studies were conducted in accordance with the Declaration of Helsinki.

### Anti-hPF4 moAbs studied

We generated and studied three different anti-hPF4 moAbs, their Fc-modified moAbs, and an isotype control moAb. The anti-hPF4 moAbs included RTO, a murine anti-hPF4 IgG2bκ moAb that binds to hPF4^19^ and prevents hPF4 monomeric units from assembling into dimeric and tetrameric hPF4^15^. RTO was used without modification. 1E12 is a chimeric anti-hPF4 moAb that binds better when hPF4 is not complexed to a polyanion^20^. 1E12 targets the VITT antigenic site of hPF4, which overlaps with the heparin-binding domain on the hPF4 tetramer^21^. Deglycosylated 1E12 (DG-1E12) was studied as well, as it no longer Fc-activates platelets^22^. KKO is a murine IgG2bκ moAb that binds hPF4 with greater affinity when the chemokine is complexed to a polyanion and does so distal to the heparin-binding domain. It does not bind mPF4^19^. G4KKO is an Fc-modified KKO that has the Fc portion of KKO switched with that of human IgG4 as detailed in **Figure *S*1**. F(ab’)2-fragments from G4KKO (FabG4KKO) were prepared using FabRICATOR Fab2 Kit (Genovis) as per the manufacturer^23^. TRA is an isotype control IgG2bκ moAb for KKO and RTO^19^ and was used unmodified. KKO, TRA and RTO were purified from hybridoma supernatants^19^.

### DLS studies of β2GPI, hPF4, DNA and APS antibodies

Studies of complex formation between β2GPI, hPF4, and genomic DNA with APS IgGs were performed as described previously for HIT^24^. β2GPI was isolated from fresh frozen plasma^25^. Briefly, plasma was treated with perchloric acid precipitation, followed by affinity chromatography using a HiTrap Heparin column (Cytiva) and ion exchange chromatography on a Mono S column (Cytiva). Fractions containing a single homogeneous band at ∼50□Da on SDS-PAGE were pooled and buffer-exchanged into physiologic buffered solution (Gibco) using ultrafiltration with 10□Da cutoff filters (Amicon Ultra, Millipore). Recombinant hPF4 was generated, purified, resuspended in 20 mM Hepes in 0.5-1 M NaCl and characterized as we described^26^. Stock solutions of β2GPI, hPF4, calf thymus DNA (Sigma), and antibodies were spun at 21,000g for 10 minutes at 4°C immediately prior to use. hPF4 was confirmed to be <10 nm by DLS prior to study. Immune complexes were formed by incubating β2GPI (10 µg/ml), hPF4 (5 µg/ml), DNA (0.5 µg/ml) and the APS IgG of interest or control IgG (300-500 µg/ml) for 5 minutes at room temperature (RT) in Dulbecco’s-PBS (DPBS-Ca^+2^/Mg^+2^: NaCl 136 mM, CaCl_2_ 0.901 mM, KCl 2.68 mM, MgCl_2_ 0.491 mM, and Na_2_HPO_4_ 15 mM) that was passed through a 0.22 μm pore filter (MilliporeSigma). To assess disruption of complex formation by the hPF4-targeting moAbs, RTO, 1E12 and G4KKO or control moAb TRA or HIgG4 as a control for G4KKO (HyHEL-10, Invitrogen), each, at 0-100 µg/ml, were complexed with hPF4 for 10 minutes before adding the other reactants to study association or were added 30 minutes post-complex-formation to study dissociation. Particle sizes were analyzed by DLS on a fixed scattering angle Zetasizer Nano-ZS system (Malvern Instruments Ltd)^20,27,28^. The Z-average size distribution (i.e., hydrodynamic diameters) of particles based on volume, was measured at RT with light backscattering of 173°. At least 25 repetitive measurements were made at the studied time points. Data analysis was performed using the Zetasizer software, version 7.03 (Malvern Instruments Ltd).

### Microfluidic channel studies of NETs

Binding studies to NETs of β2GPI and/or hPF4 were conducted in a BioFlux microfluidic system (Fluxion Biosciences) as we described^16^ using human neutrophils isolated by lympholyte density gradient (Cedarlane). Isolated neutrophils (2 x 10^6^ cells/ml) were incubated with 1 ng/ml tumor necrosis factor-α (Gibco) at RT for 10 minutes and then flowed through channels coated with fibronectin (Sigma) resulting in neutrophil adhesion. The cells were then exposed to 100 ng/ml phorbol-myristyl-acetate (PMA) overnight at 37°C. Released NETs were visualized with 1 μM sytox-orange (ThermoFisher). The channels were infused with hPF4 (0-25 μg/ml) and β2GPI (0-20 μg/ml) for 30 minutes. Binding of β2GPI was measured using an Axio Observer Z1 inverted microscope (Zeiss) equipped with a motorized stage and an HXP-120 C metal halide illumination source by infusion of Alexa Fluor488-labeled mouse anti-β2GPI moAb (50 µg/ml, cat no: NBP3-26956AF488, Novus Biologicals). The microscope and image acquisition were controlled by BioFlux Montage software with a MetaMorph-based platform (Molecular Devices). Z-stacks of β2GPI binding to the NETs were captured using a Zeiss LSM 710 Confocal microscope. The final data (mean Z-stack) were plotted and analyzed using ImageJ.

DNase1-resistance of NET-lined microfluidic channels was analyzed in the absence and presence of hPF4 (6.5 µg/ml) and β2GPI (20 µg/ml), similar to our prior studies with hPF4^29^. These studies utilized the same microfluidic system to examine β2GPI binding to NETs, but with the addition of patient-derived APS IgG (100 µg/ml). Subsequently, DNase1 (100 U/ml, Sigma-Aldrich) was added to the system for 10 minutes, and remaining sytox-orange staining was detected.

### Microfluidic channel studies of thrombosis

*In vitro* thrombosis studies under flow were performed in the BioFlux device with the channels coated with near-confluent HUVECs (Lifeline Cell Technology)^30^. HUVECs were injured by infusion of hematoporphyrin (50 µg/ml, Sigma) while exposing the cells for 20 seconds to an HXP120C light source (Zeiss) with a 475nm excitation and 530nm emission filter. Whole blood from healthy controls in sodium citrate (final concentration of 0.32%) was infused through these channels. Prior to infusion, some blood samples were incubated for 15 minutes with hPF4 (25 μg/ml) and/or β2GPI (20 µg/ml) along without added IgG or with 100 µg/ml IgG isolated from patients with APS or healthy controls. To detect platelet thrombus and fibrin generation on the injured HUVECs from onset of infusion until 15 minutes after starting the infusion, blood samples were preincubated with an anti-human-CD41-Alexa Fluor488 moAb (#11-0419-42; Invitrogen) and anti-human-fibrin Alexa Fluor561 moAb^31^. In some experiments, either DNase1 (100 µg/ml) or ADAMTS13 (0.7 mg/ml, R&D Systems) were added to the whole blood 15 minutes prior to infusion into the channels. In others, the anti-hPF4 mo Abs RTO, 1E12, or KKO or their Fc-modified versions or the control moAb TRA (each at 100 µg/ml) were added to the samples.

To examine complement activation, HUVEC nuclei were stained after completing the 15-minute infusion using Hoechst (1:5000 dilution, ThermoFisher) and accumulated C1q was stained using an Alexa Fluor594-labeled murine anti-human C1q moAb (#sc-53544; Santa Cruz) and C5b-9 using an Alexa Fluor647-labeled murine anti-C5b-9 moAb (#NBP1-05120AF-647: Novus Biological). Z-stack images of injured endothelium were captured using a Zeiss LSM 710 Confocal microscope. The final data (mean Z-stack) was plotted and analyzed using ImageJ.

### Passive immunization APS murine model

All murine studies were done in 8-12-week-old C57BL/6 wildtype (WT) or mice that were littermates that either were knockout mPF4 (termed mPF4^-/-^) or transgenic hPF4^+^/mPF4^-/-^ (termed hPF4^+^) or hPF4^+^/FcγRIIA^+^/mPF4^-/-^ (termed HIT) mice. Institutional animal care and use committee approved the murine studies, and animals were euthanized in accordance with the American Veterinary Medical Association.

Neutrophil rolling and adhesion to cremaster arterioles and venules were studied. Mice were anesthetized with intraperitoneally injected pentobarbital sodium (11 µg/g, Abbott), had their cremaster arterioles and venules exposed, and their intraluminal space visualized using confocal intravital microscopy^16^. Neutrophils and neutrophil extracellular vesicles (EVs, defined by diameter ≤ 3µ)^32^ were labeled by infusing intrajugular (IJ) F(ab’)2-fragments of Alexa Fluor 488-labeled anti-murine Ly6G antibody (0.3 µg/g mice, clone 1A8; BD Bioscience). Studies were done pre- and post-infusion of APS1 or control IgGs (10 µg/g, IJ) in WT, hPF4^+^, HIT, and mPF4^-/-^ mice. hPF4^+^ mice were also studied using APS2 or control IgG. Confocal time-lapse images of neutrophil rolling and adherence were collected using Slidebook 6.0 (Intelligent Imaging Innovations) and analyzed for neutrophil rolling speed using Imaris 10.2 (Oxford Instruments). In some of the APS2 moAb infusion studies, either G4KKO or RTO or TRA moAb (each at 2 µg/g) were infused as a “prophylaxis” therapy 15 minutes prior to APS2 or control IgG, or “therapeutically” 15 minutes after APS2 or control IgG.

For cremaster arteriole and venule laser injury studies, the same genotypic mice were prepared as described for the neutrophil rolling and adhesion studies^16^. Laser injuries were induced using an SRS NL100 Nitrogen Laser system (Photonic Instruments). Platelets were fluorescently labeled with Alexa Fluor647-anti-CD41 F(ab′)_2_ fragments (0.3□µg/g mouse, IJ, BD Bioscience), and fibrin with Alexa Fluor561-anti-fibrin antibody (0.3□µg/g mouse, IJ)^16^. Neutrophils and neutrophil extracellular vesicles (EVs, <3µ in diameter) incorporated within the thrombi were identified by the binding of Alexa Fluor488-labeled Ly6G antibody. Data collection and confocal time-lapsed images of the thrombi were collected over 3 minutes post injury and analyzed using Slidebook 6.0 (Intelligent Imaging Innovations) to identify the volume of the platelet and fibrin thrombi and to enumerate the number of incorporated neutrophils and neutrophil EVs per clot.

### Statistical analysis

Differences between 2 groups were compared using a two-tailed Student t-test. For multiple comparisons of more than 2 groups, statistical analysis was performed by two-way analysis of variance (ANOVA) using GraphPad Prism version 10.42. Differences were considered significant when P < 0.05.

## Results

### Formation of β2GPI:hPF4:DNA APS antigenic complexes

DLS can be used to study protein-protein interactions in solution, avoiding non-specific interactions with plastic surfaces^24^. We first asked if hPF4 forms complexes with β2GPI in the absence or presence of DNA, the polyanionic backbone of NETs^33^. We confirmed the prior observations that β2GPI complexes with hPF4^11^ and that each can complex with DNA (**Figure 1A**). As expected, hPF4, which is known to aggregate DNA^16,29^, formed larger complexes with DNA than did β2GPI^18^. The largest molecular weight complexes were formed when all three were present (hPF4:β2GPI:DNA, **Figures 1A** and ***S*2**). APS1 and APS2 IgGs did not form large complexes with hPF4:β2GPI or β2GPI:DNA, but did so with hPF4:β2GPI:DNA (**Figures 1B** and ***S*2**).

**Figure 1.**
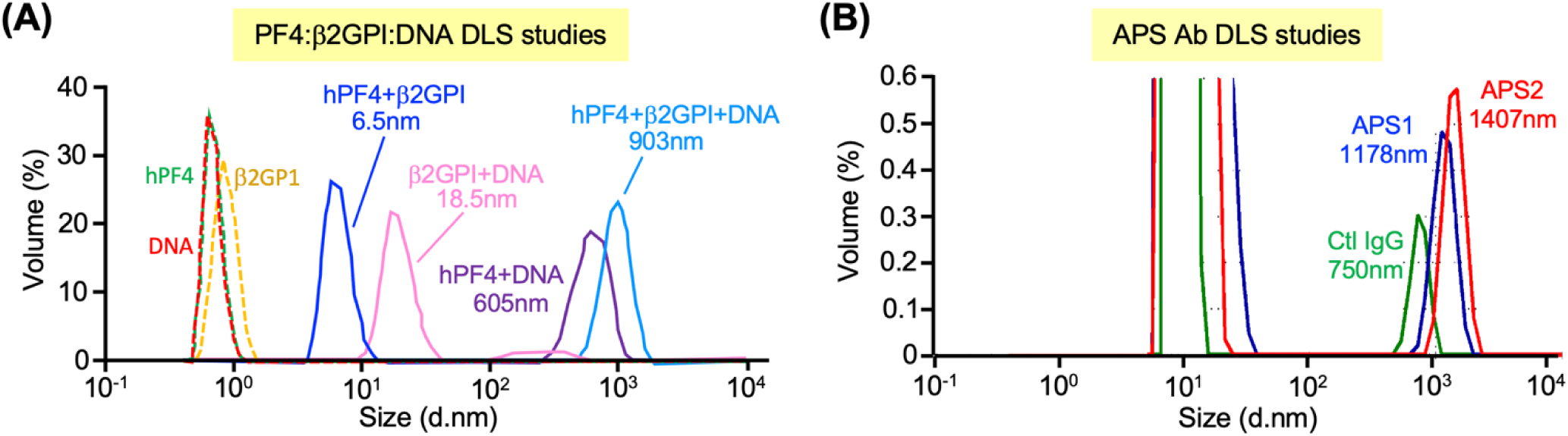
DLS studies in the presence of β2GPI, hPF4, DNA and APS antibodies. Representative DLS studies done with N ≥3 experiments per arm. Histogram showing particle sizes based on volume. (**A**) Studies of β2GPI (10 µg/ml), hPF4 (5 µg/ml) or calf thymus DNA (0.5 µg/ml) alone or in combination. Complex formation at room temperature was analyzed by DLS at 20 hours. (**B**) DLS studies of β2GPI:hPF4:DNA in the presence of added IgG from two different patients with APS or from a healthy control (each at 500 µg/ml). See also **Figure *S*2** for DLS studies with APS2 antibody complexed with either β2GPI:DNA or hPF4:β2GPI or β2GPI:hPF4:DNA.

### Microfluidic studies of β2GPI binding to NETs: the role of hPF4

To further investigate binding of β2GPI to NETs, we studied binding in a previously described PMA-activated, neutrophil-released, NET-lined microfluidic system^16,29^. Unlike in the DLS studies, this system is amenable to venous flow rates and can be performed in the complex environment of whole blood^31^. In this system, β2GPI alone bound to NETs poorly (**Figures 2A** and **2B**). The addition of hPF4 at a concentration that can be reached at sites of thrombosis (20 µg/ml)^34^, resulted in an approximately seven-fold increase in β2GPI binding over background (**Figures 2A** and **2B**). In the presence of APS antibodies, binding of β2GPI to NETs increased by approximately five-fold by inclusion of hPF4 (**Figures 2A** and **2B**).

**Figure 2.**
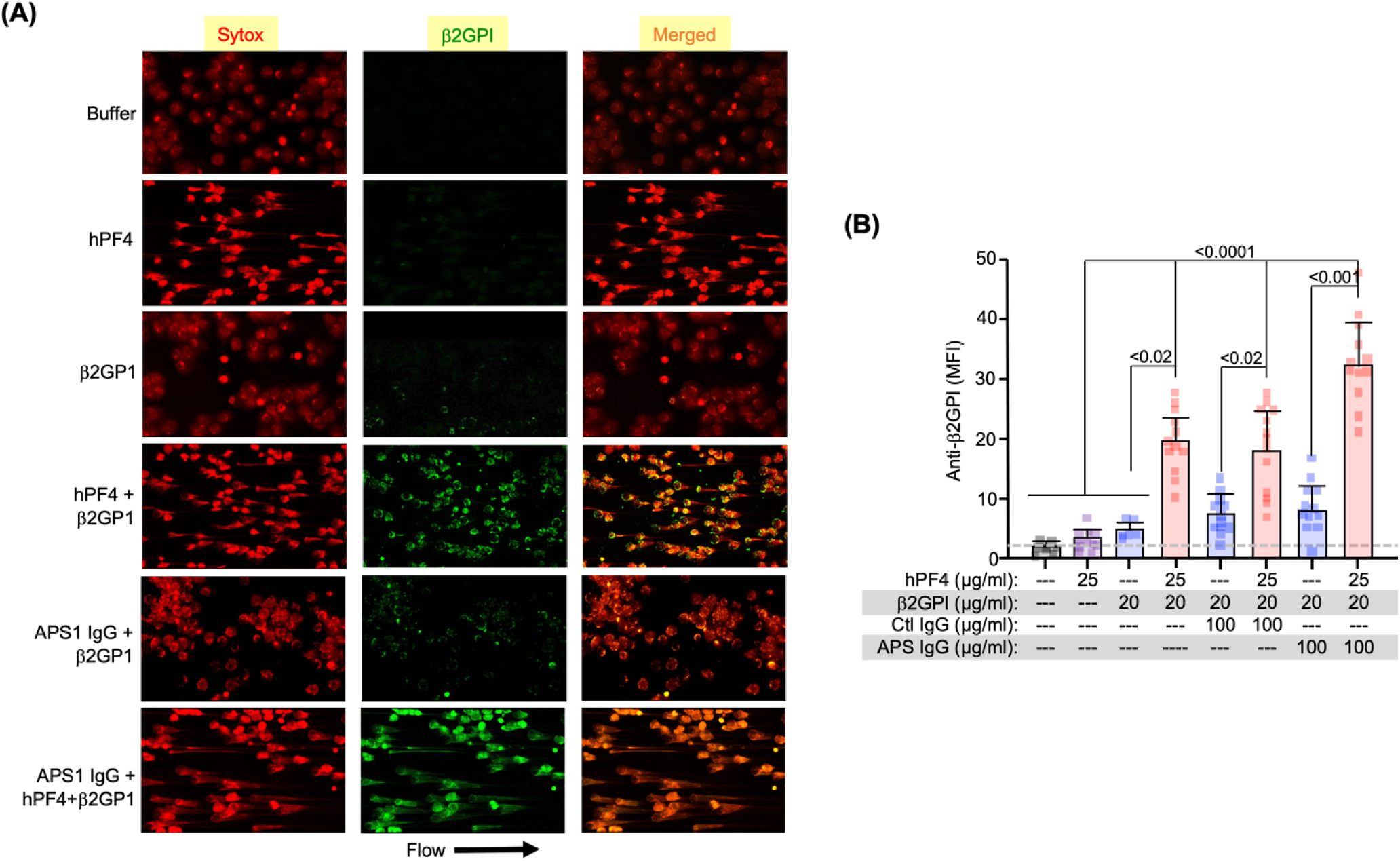
NET microfluidic studies of β2GPI in the presence of hPF4 and APS antibodies. **(A)** Representative microfluidic images of PMA-activated neutrophils and released NETs in the absence and presence of 25 µg/ml of hPF4 and/or 20 µg/ml β2GPI. Studies shown were done with APS1 IgG at 100 µg/ml. Sytox-orange staining of released NETs (red) and β2GPI detected using a polyclonal anti-β2GPI (green). Arrow indicates direction of flow. (**B**) Same as in (A), but aggregate of studies with an equal number of studies with APS1 and APS2 antibodies. Shown are mean fluorescent intensity (MFI) for β2GPI bound to the released NETs ± 1 standard deviation (SD) for N = 12 per arm, measured in duplicate. Individual data points are shown. Grey dashed line represents background anti-β2GPI antibody binding to the channel. Differences were examined by two-way ANOVA.

We previously showed that KKO and HIT IgGs can protect PF4:NET complexes from DNase1 digestion, likely by sterically blocking the enzyme from attacking the phosphoribose backbone of the DNA^16^, thereby stabilizing the hPF4:NET:HIT antibody complexes and contributing to the prothrombotic nature of HIT. We found that hPF4:β2GPI:NET:APS IgG immune complexes also were resistant to lysis by DNase1 in this microfluidic system compared to hPF4:β2GPI:NETs:control IgG (p < 0.005, **Figure 3A**).

**Figure 3.**
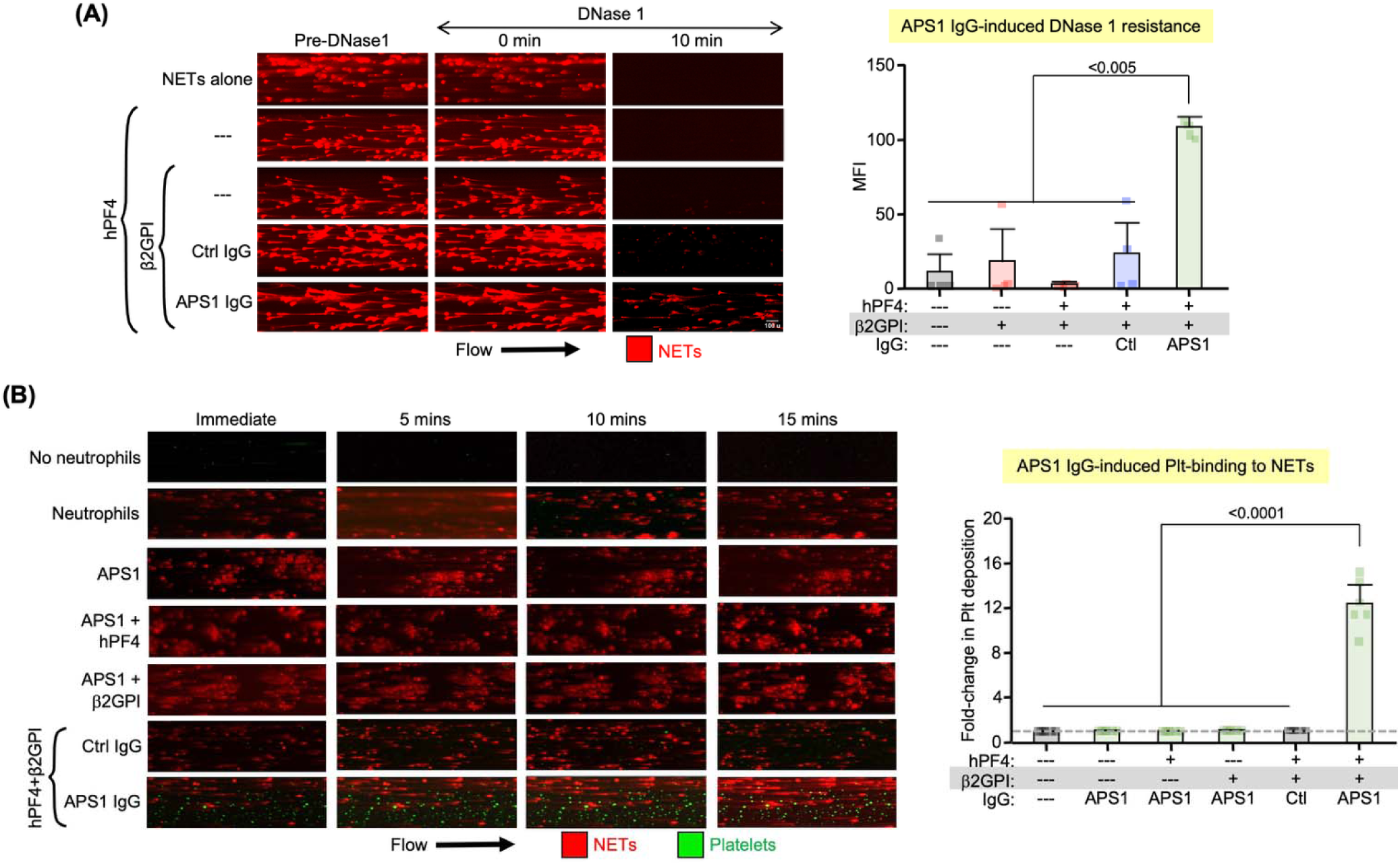
NET-lined microfluidic studies. **(A)** <u>Left</u>: Representative images from studies of DNase1 resistance of sytox-orange stained extruded chromatin-lined channels from PMA-activated neutrophils. Channels supplemented with hPF4 and/or β2GPI with or without APS1 or healthy control IgG at the same concentrations as in Figure 2. <u>Right</u>: Mean ± 1 SD for N = 4 studies, each done in triplicate for remaining sytox-orange-stained DNA after DNase1. Statistical analysis is shown for arm with added APS1 compared to the other arms by two-way ANOVA. (**B**) <u>Left</u>: Representative images from studies of channels lined with neutrophils in the presence of sytox-orange. Healthy whole blood with platelets labeled with an anti-human-CD41-Alexa Fluor488 moAb was flowed through channel supplemented as in Figure 2. <u>Right</u>: Mean ± 1 SD for relative bound platelets to channels lined with NETs extruded from adherent neutrophils, but no added hPF4 or β2GPI or IgG (represented by dashed, grey line). N = 6 studies, each done in triplicate. Statistical analysis is shown for studies comparing platelet binding in the presence of control (Ctl) IgG to APS1 IgG by one-way ANOVA. See bar graph analysis for similar studies using APS2 or APS3 IgGs in **Figure *S*3**.

We then studied microfluidic channels lined with neutrophils stimulated to release NETs. Whole blood from healthy controls was flowed through the channels in the absence or presence of added β2GPI and hPF4 along with either control or APS IgG (**Figure 3B**). Platelet adhesion to the NETs was measured. In this model system, platelet adhesion to the NETs was not observed when APS1 IgG was added alone or in combination with either hPF4 or β2GPI, but did so when hPF4 plus β2GPI was added (**Figures 3B**), consistent with the known ability of β2GPI-positive APS antibodies to interact with neutrophils and NETs and be prothrombotic^17,18^. Similarly, platelet thrombi formed on NETs in the presence of APS2 and APS3 IgGs, also only in the presence of both hPF4 and β2GPI (**Figure *S*3**).

### Endothelialized microfluidic studies of PF4 in APS antibody-induced thrombosis

We previously developed a photochemically-injured, HUVEC-lined, microfluidic system to study the mechanistic basis of HIT using whole blood from healthy donors to which HIT-like moAbs or IgG from patients with HIT was added^31^. Similar studies were now done with adding APS IgGs as depicted in the schema in **Figure *S*4**. We found that platelet-rich thrombi and fibrin accumulation were minimal unless both hPF4 and β2GPI were added to the blood sample (**Figures 4A** and ***S*5** for platelets, and **Figure *S*6** for fibrinogen). The results were consistent with all four APS IgG preparations tested (**Figures 4A** and ***S*5**). We also stained the channels after thrombus formation for C1q and C5b-9 deposition as indicators of complement activation^35^, and found that complement deposition was seen only when all three – β2GPI, hPF4 and APS IgG – were added to the blood samples (**Figures 4B** for C5b-9 and C1q).

**Figure 4.**
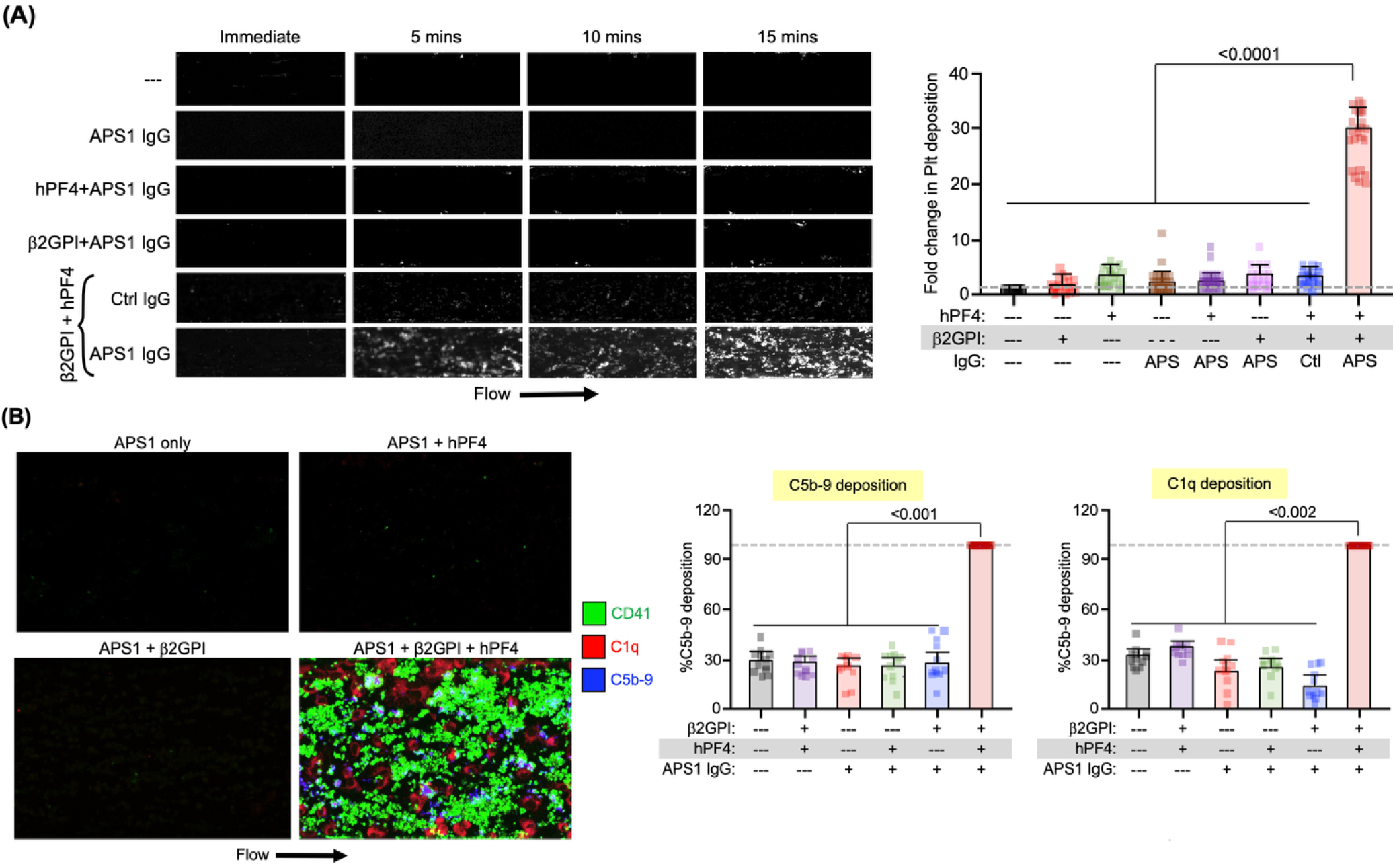
Photochemically injured HUVEC-lined microfluidic studies. See **Figure *S*4** for a schematic of the injured HUVEC microfluidic system. (**A**) <u>Left</u>: Representative images from whole blood flowed through uninjured and injured HUVEC channels. The injured channels were supplemented with hPF4 (25 µg/ml) and/or β2GPI (20 µg/ml) and/or APS1 IgG (50 µg/ml). <u>Right</u>: Mean ± 1 SD of platelet (Plt) deposition in the presence of added hPF4 and/or β2GPI and/or IgG from APS1 through APS4 and control (Ctl) compared to nothing added as indicated by the dashed grey line. N = 4 healthy donors studies, each study was done in triplicate and then aggregated for APS1 through APS4 IgG studies. Statistical analysis is shown for study with added hPF4 plus β2GPI plus APS IgGs compared to the other studies by two-way ANOVA. See **Figure *S*5** for representative studies with the individual APS2 through APS4 IgGs, and **Figure *S*6** for overall fibrin deposition. (**B**) Same study as in (A), but post-facto staining for C5b-9 and C1q as well as platelets. <u>Left</u>: Representative image post-completion of the study in (A) stained for C5b-9 and C1q in the study done under indicated conditions. <u>Middle</u>: Mean ± 1 SD of C5b-9 deposition in the presence of hPF4 plus β2GPI plus APS IgGs compared to other conditions studied. N = 12 per arm. Statistical analysis is shown for studies done with added APS IgG compared to the other studies by one-way ANOVA. <u>Right</u>: Similar study, but for C1q deposition.

We previously showed that thrombus formation in the photochemically-injured HUVEC model depends on the release of von Willebrand factor (VWF) from the injured HUVEC lining^36^. We now asked whether VWF or NETs were involved in thrombosis induced by APS IgG (**Figure 4A)**. When the studies in **Figure 4A** were repeated including DNase 1 or ADAMTS13, we found that both contributed to the development of platelet thrombi (**Figure *S*7**).

### Venous neutrophil rolling post-APS IgG infusion in a passive-infusion murine model of APS

We previously reported that in hPF4^+^/FcγRIIA^+^/mPF4^-/-^, termed HIT, mice, neutrophil-rolling on uninjured venules slowed significantly almost immediately post-infusion of HIT antibodies accompanied by increased neutrophil adherence to the endothelial lining^16^. This increase in neutrophil rolling and adherence to venules was associated with greater accumulation of neutrophils surrounding and within a subsequent cremaster venule laser injury-induced thrombus, enhancing thrombosis^16^. We now observed the same change in neutrophil:venule wall interaction in WT, hPF4^+^ and HIT mice after infusion of IgG isolated from a patient with APS into uninjured cremaster venules (**Figures 5A** and **5B**; see also **Videos 1** through **4** for pre- and post-infusion of APS1 IgG into WT and hPF4^+^ mice, respectively). In contrast, infusion of APS1 antibodies into mPF4^-/-^ mice did not alter neutrophil rolling or adhesion compared to pre-transfusion (**Figures 5A** and **5B**; see also **Videos 5** and **6** for pre- and post-infusion of APS1 IgG into mPF4^-/-^ mice). Parallel studies with control IgG did not alter neutrophil rolling and adhesion to the endothelium in any mice genotype (**Figure *S*8**).

**Figure 5.**
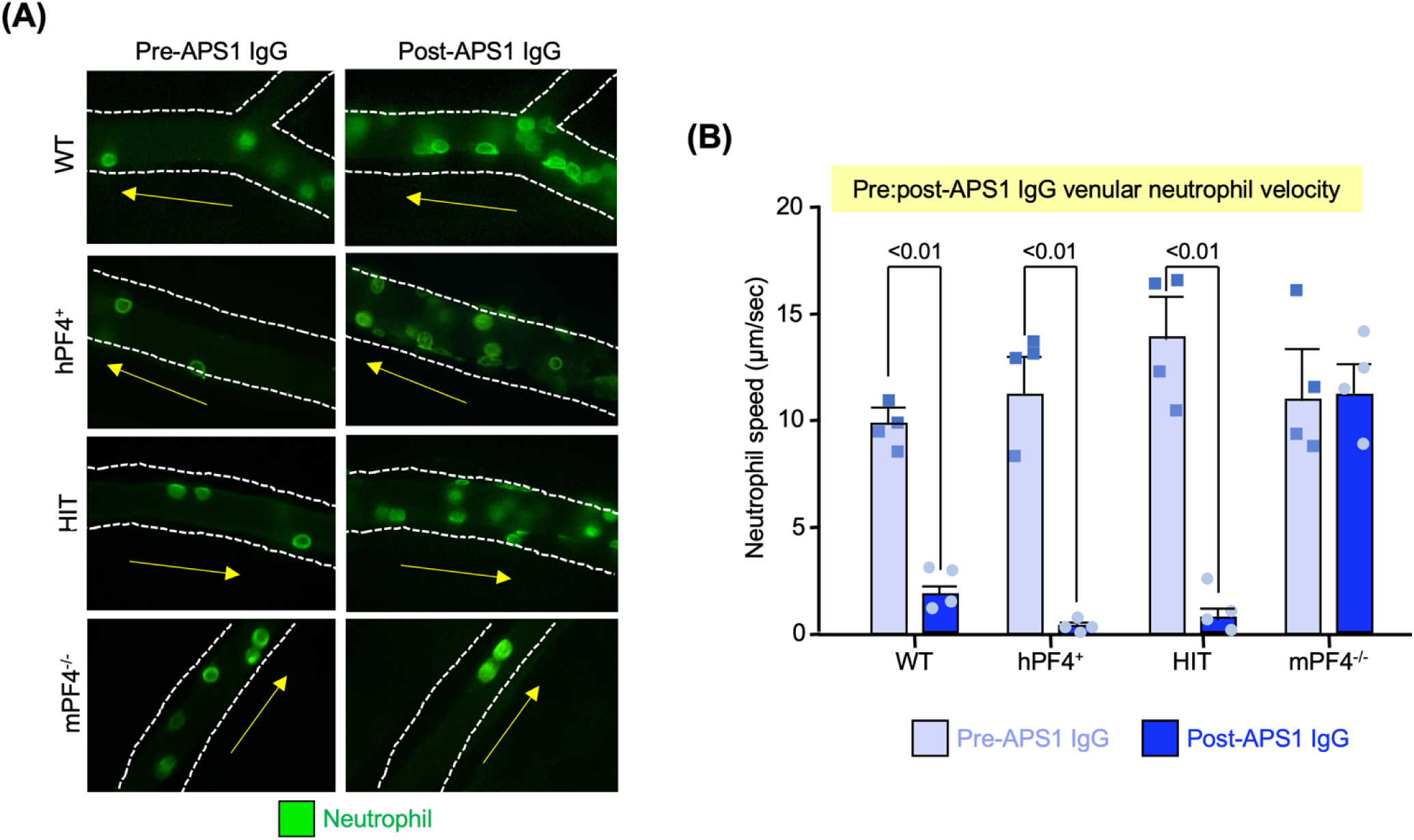
Neutrophil rolling on cremaster venules. **(A)** Representative images of neutrophil venule rolling looking at the identical vessel pre- and 10 minutes post-APS1 IgG given IV. Studies were done in 4 different genotype mice: WT, hPF4, HIT and mPF4^-/-^ mice. The direction of venous flow is indicated by a yellow arrow and the vessel walls outlined by a white dashed line. (**B**) Mean ± 1 SD for neutrophil rolling speed both pre- and post-APS1 IgG infusion. N = 4 mice per genotype. P values determined by two-way Student t-test to compare pre- *vs*. post-APS IgG infusion. See **Figure *S*8** for similar studies done with control IgG.

### Arterial and venule thrombi post-APS IgG infusion in a passive-infusion model of APS in mice

Cremaster venule and arteriole thrombi were induced by laser injury as we described in murine studies of HIT^16^, were now conducted in WT, hPF4^+^, HIT and mPF4^-/-^ mice using APS1 IgG and healthy control IgG. Laser-induced venule thrombi were enlarged in WT, hPF4^+^ and HIT mice receiving APS1 as measured by platelet and fibrin incorporation compared to littermate mPF4^-/-^ mice and to mice that had received control IgG (**Figures 6A** for APS1 infused mice and ***S*9A** for control IgG infused mice; see **Video 7** for APS1 IgG-enhanced thrombosis in HIT mice compared to **Video 8** for a similar study in mPF4^-/-^ mice). APS1 IgG infusion also enhanced laser-induced arterial thrombus growth as measured by platelet and fibrin accumulation in WT, hPF4^+^ and HIT mice compared to mPF4^-/-^ mice and to mice that had received control IgG (**Figures 6B** and ***S*9B**; see also **Video 9** for HIT mice and **Video 10** for mPF4^-/-^ mice receiving APS1 IgG).

**Figure 6.**
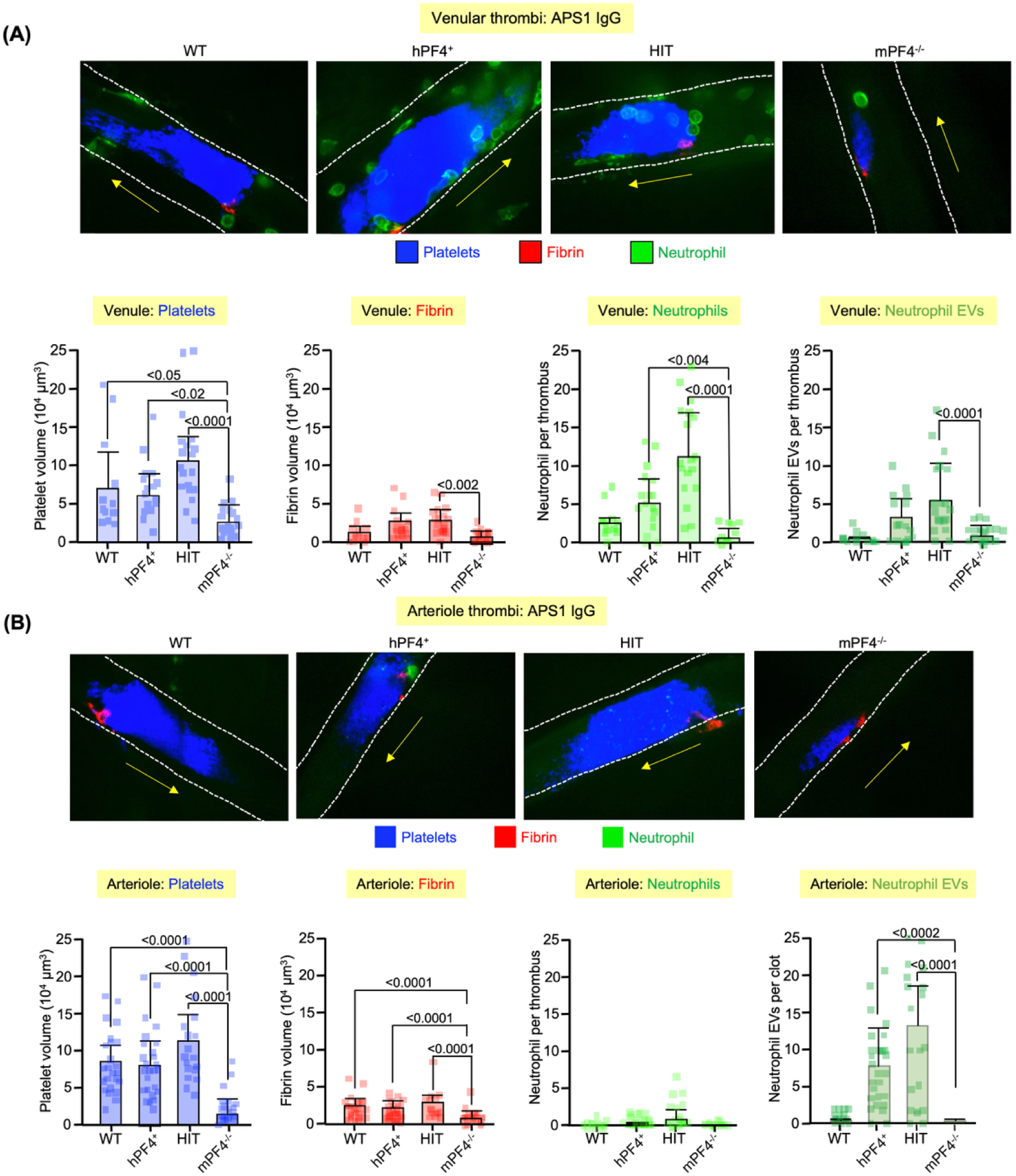
Thrombus formation in a passive inoculation APS murine model. Similar to the studies in Figure 5, but a laser injury is induced to the cremaster venule or arteriole, and incorporation of platelets, fibrin, neutrophils and neutrophil EVs followed. (**A**) Venule injuries were performed. <u>Top row</u>: Representative thrombi. The yellow arrow indicates direction of blood flow and the dashed line outlines the vessels. <u>Bottom row</u>: Graphs of mean ± 1 SD for platelet, fibrin, neutrophil and neutrophil EV incorporation into thrombi as well as individual data points are shown. For each arm, N = 3-5 mice and 8-20 injuries done in aggregate for each arm. Statistical analysis was by one-way ANOVA. Statistical comparisons shown are with the mPF4^-/-^ mice compared to the other genotypic mice groups. (**B**) Similar studies to (A), but for arterioles. Here, there were N = 5-9 mice per arm with 19-29 injuries done in aggregate for each arm. See **Figure *S*9** for parallel studies with healthy control IgG studied instead of APS1.

Neutrophil incorporation into the venule and arterial thrombi was also observed (**Figure 6**). Intact neutrophils were incorporated into the venule thrombi in the APS1-treated mice, with the greatest incorporation occurring in the HIT mice followed by the hPF4^+^ mice with the least being in the mPF4^-/-^ mice (**Figure 6A**). Mice infused with control IgG showed few neutrophils incorporated into venule thrombi independent of genotype (**Figure *S*9A**). Arterial thrombi in the APS1-treated mice had few neutrophils, but more neutrophil EVs (**Figure 6B**), with the highest level in the HIT mice followed by the hPF4^+^ mice, while the mPF4^-/-^ mice had levels near that seen in control IgG infusion studies in any mice (**Figure *S*9B**).

### Anti-hPF4 moAbs intervention in APS pathobiology: in vitro studies

We next examined whether any of three anti-hPF4 moAbs, RTO, 1E12 or KKO or their Fc-modified versions, can block these immune complexes from forming and potentially serve as alternative therapeutics to anticoagulants for patients with anti-β2GPI-positive APS. DLS studies showed that RTO and 1E12 prevented the formation of large hPF4:β2GPI:DNA immune complexes with APS1, resulting in major peaks at much smaller size, representing either individual components or small complexes (**Figure 7A**). An Fc-modified KKO on an IgG4 subclass, termed G4KKO, and characterized in **Figure *S*1** increased the size of the observed complexes, likely crosslinking the APS immune complexes via hPF4 (**Figure 7A**).

**Figure 7.**
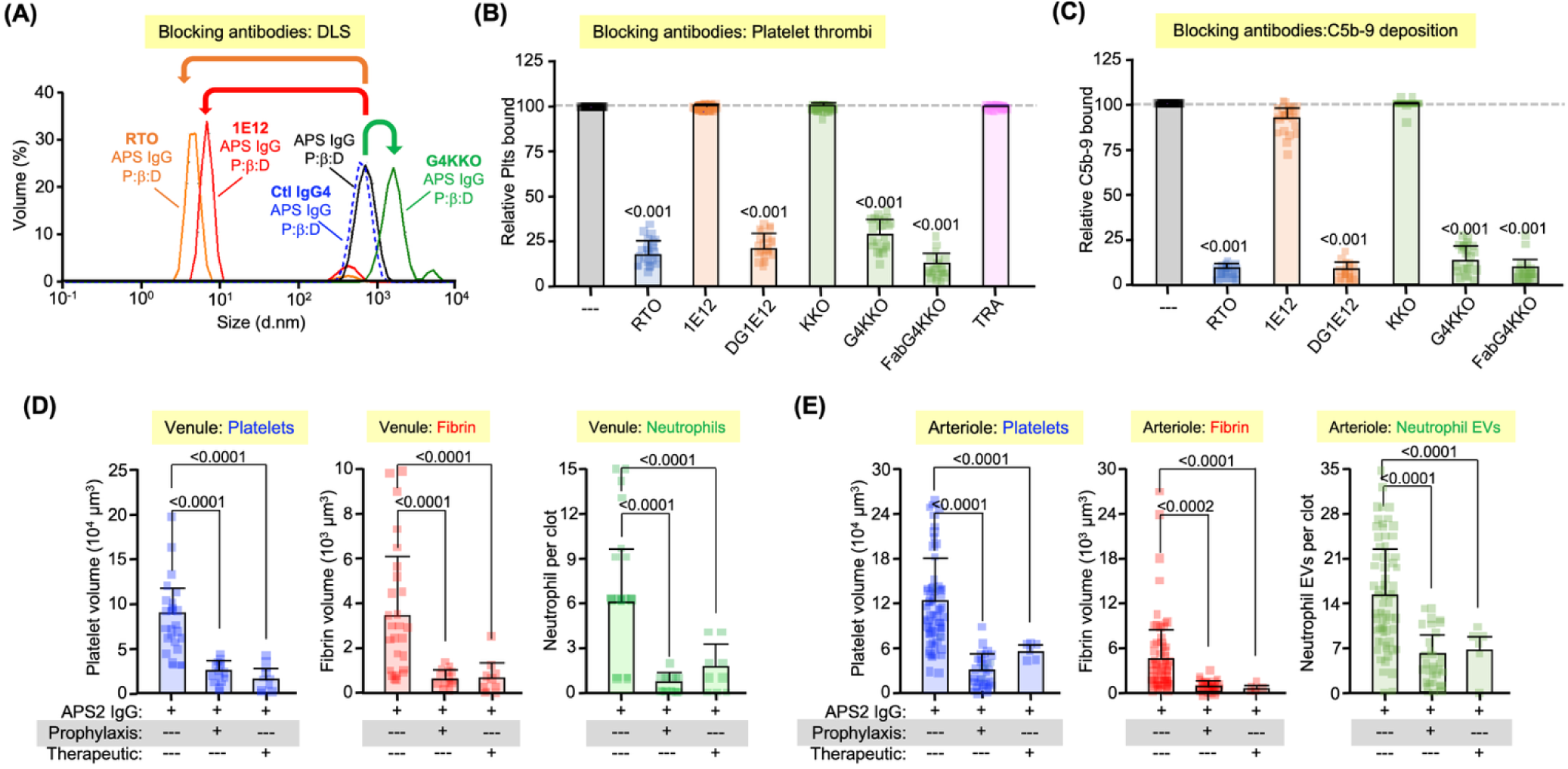
Use of anti-hPF4 moAbs in APS to block immune complex formation and thrombosis. **(A)** DLS studies as in Figure 1 in the presence of hPF4 plus β2GPI plus DNA (black). The addition of RTO (red, 100 µg/ml) or 1E12 (orange, 35 µg/ml), resulted in loss of the antigenic complex large peak at 10^2-3^ nm and the appearance of small peaks at ∼10^-1^ nm. G4KKO (35 µg/ml) resulted in a larger complex of > 10^3^ nm. Data shown are representative of 3 independent experiments performed at 2 hours post incubation. (**B**) Injured HUVEC microfluidic studies of platelet incorporation into thrombi as in Figure 4 and the effects of the various anti-hPF4 moAbs and TRA, an isotype control moAb for KKO and RTO (each at 100 µg/ml). Mean ± 1 SD of platelets bound in the presence of the antibody compared to no blocking antibody (grey bar and dashed grey line). N = 24 studies in aggregate of APS1 through APS3, each measured in duplicate, with blood from 4 separate healthy donors. Individual data points are shown. P values were determined by two-way ANOVA. P values shown are those that significantly differed from no added anti-hPF4 moAb. See **Figure *S*10** for representative images of the tested anti-hPF4 moAbs for each individual tested anti-hPF4 moAb and mean ± 1 SD analysis. See **Figure *S*11** for fibrin incorporation. (**C**) Bar graph as in (B), but for C5b-9 incorporation. See **Figure *S*12** for representative images and mean ± 1 SD analysis for C1q deposition. (**D**) and (**E**) The effects of G4KKO infused IV (2 µg/g) to hPF4^+^ mice treated with 10 µg/g, IJ, APS2 IgG, on platelet, fibrin and neutrophil incorporation into venule thrombi in venules and neutrophil EVs in arterioles, respectively. In (D), N = 3-7 mice per arm. In (E), N = 3-7 mice per arm. P values were determined by two-way ANOVA. Only significant comparative differences between those receiving or not receiving G4KKO are shown. See **Figure *S*13** for representative images and quantitation of G4KKO and RTO effects on neutrophil rolling in the same APS2 IgG/hPF4^+^ mice model given either prophylactically or therapeutically. **Figure *S*14** shows representative venule and arteriole thrombi after APS2 IgG infusion in hPF4^+^ mice with G4KKO and RTO given either prophylactic or therapeutic. **Figure *S*15** provides the quantitative analysis of effects of APS2 IgG-enhanced thrombosis in the presence of either prophylactic or therapeutic RTO.

Microfluidic studies on injured HUVEC were conducted in the presence of modified anti-hPF4 moAbs. RTO and the isotype control TRA were studied unmodified. 1E12 was studied unmodified and after deglycosylation to prevent FcγRIIA and complement activation (DG-1E12)^22^. KKO was studied unmodified and as G4KKO. G4KKO was used before and after proteolysis into a F(ab’_2_) fragment (FabG4KKO). Unmodified KKO or 1E12 and also TRA as a negative control did not decrease the size of thrombi (**Figure 7B**). In contrast, RTO, DG-1E12, G4KKO and FabG4KKO all decreased platelet thrombus size (**Figures 7B**) irrespective of whether APS1 or APS2 or APS3 IgG was studied (**Figures *S*10A** through ***S*10C**). Additionally, these moAbs also decreased fibrin clot formation (**Figure *S*11**), and C5b-9 and C1q deposition (**Figures 7C** and ***S*12**, respectively).

### Anti-hPF4 moAbs intervention in APS pathobiology: in vivo studies

Studies of anti-hPF4 moAbs blocking the prothrombotic effects of APS IgG in murine models focused on G4KKO and RTO intervention after infusion of APS2 IgG into hPF4^+^ mice. We studied APS2 IgG to demonstrate *in vivo* studies with a second APS IgG preparation and using hPF4^+^ mice. **Figures 7D**, **7E**, ***S*13** and ***S*14** gave similar results with those seen with APS1 IgG in WT, hPF4^+^ and HIT mice (**Figure 6**). The data in **Figure *S*1B** show that G4KKO bound similarly to PF4:heparin as KKO. While G4KKO showed reduced, but not absent, activation of FcγRIIA^+^ human and mouse platelets, it does not activate hPF4^+^ mouse platelets as they lack this Fc receptor (**Figures *S*1C** and ***S*1D**, respectively). Also, unlike KKO, G4KKO does not activate complement (**Figure *S*1E**). Neutrophil rolling studies showed that both G4KKO and RTO were effective in limiting the effects of APS2 IgG infusion on rolling in HIT mice when given 15 minutes prior to infusing the APS2 IgG (**Figure *S*13A**), but caused less improvement in neutrophil rolling when given 15 minutes post, especially for RTO (**Figure *S*13B**).

The effect of G4KKO and RTO on APS-enhanced thrombosis were examined using laser-induced cremaster venule and arteriole injuries in hPF4^+^ mice post-infusion of APS2 IgG (**Figures 7D**, ***S*14A** and ***S*15A**, and **Figures 7E**, ***S*14B** and ***S*15B**, respectively for venule and arteriole). APS2 IgG alone enhanced both arteriole and venule thrombosis in hPF4^+^ mice compared to control IgG (**Figure 7D** and ***S*15A** for venous thrombi and **Figure 7E** and ***S*15B** for arterial thrombi). In venule and arteriole injuries, G4KKO and RTO, whether again given prophylactically or therapeutically. Both reduced platelet thrombus mass and fibrin accumulation due to APS2 IgG infusion (**Figure 7D** and ***S*15A** for venule injuries, and **Figure 7E** and ***S*15B** for arteriole injuries).

G4KKO reduced the accumulated neutrophil mass in venule thrombi when given both prophylactically, or therapeutically (**Figures 7D** and ***S*14A**). RTO had a similar effect on venous neutrophil and on neutrophil EV accumulation (**Figures S14C** and ***S*15A**). In arteriole injuries where few neutrophils, but only neutrophil EVs, are accumulated into thrombi, both G4KKO and RTO suppressed the number of neutrophil EVs in arteriole thrombi when given either prophylactically or therapeutically (**Figures 7E** and ***S*14B** for G4KKO, and ***S*14D** and ***S*15B** for RTO).

## Discussion

APS is a thromboinflammatory disorder mediated by antibodies to β2GPI in many, although not in all, patients^38^. APS can cause recurrent arterial and venous thromboemboli that necessitate indefinite anticoagulation, but this therapy predisposes to concerns about major bleeding^1–3^. In obstetrics, heparin shows incomplete effectiveness, especially in triple-positive mothers^4^. The mechanism by which antibodies to β2GPI cause thrombosis is only partially understood^39^. New mechanistic insights into the prothrombotic basis of APS may lead to novel mechanism-based, non-anticoagulant therapeutics.

Our studies are predicated, in part, on the recent finding that β2GPI binds to NETs as well as to phospholipids^18^. In two other thromboinflammatory disorders, HIT and VITT, the importance of NETs in the development of the prothrombotic state has been recognized^16,40^. PF4:NET immune complexes can lead to Fc receptor-mediated intravascular activation of platelets, neutrophils, and monocytes^13^, and activate complement^35^. Both Fc-receptor and complement activation of neutrophils enhance NET release^40^. It had also been shown that β2GPI binds to PF4, which increases binding of antibodies from patients with APS *in vitro*^11^, but the importance of PF4 in the prothrombotic state of APS and its mechanistic basis has never been reported. We now asked whether PF4 enhances β2GPI binding to NETs and whether PF4 is important in the thrombotic immunopathology of APS.

The data presented extend prior observations that showed that both β2GPI and PF4 individually bind to DNA^16,18^. We show that these three components can form higher molecular-weight complexes recognized by IgG antibodies in triple-positive APS patients, and that PF4 markedly enhances β2GPI binding to NETs and platelet binding in the presence of APS IgG. Moreover, *in vitro* studies support the concept that PF4 is needed to maximize APS thromboinflammation, including neutrophil and platelet activation, and fibrin and complement deposition. *In vivo* studies also show the role of PF4 in neutrophil adhesion to the venous lining, and platelet- and neutrophil-rich exuberant thrombosis in APS mice.

We propose that, at sites of inflammation, PF4 is released from activated platelets contemporaneously with the extrusion of NETs from activated neutrophils, forming PF4:NET complexes as in HIT^16^. PF4 stabilizes the phosphoribose backbone against circulating DNase1, leading to increased levels of PF4:NETs (**Fig. *S*16**). β2GPI binds to these stabilized PF4:NETs and attracts anti-β2GPI antibodies, forming large immune complexes that engage cell surface Fc receptors on platelets and neutrophils^41^, and activate complement via the classic pathway^35^, releasing additional NETs that further amplify the prothrombotic pathways. The need for sufficient concurrent PF4 and NET release to initiate this prothrombotic cycle may explain why patients with APS can be antibody-positive, but asymptomatic^42^.

Each of three anti-hPF4 moAbs that disrupt hPF4 biology by binding to distinct sites on hPF4 inhibited the *in vitro* and *in vivo* effects of IgG anti-β2GPI-reactive APS antibodies. We propose that the three anti-hPF4 moAbs block anti-β2GPI immunothrombosis by three distinct mechanisms (**Figure *S*16**): RTO by disrupting hPF4 tetramer formation so that the hPF4 does not bind to β2GPI or crosslink DNA within NETs; 1E12 by sterically blocking the hPF4 heparin-binding domain so that the hPF4 cannot bind to the DNA and KKO by crossbinding two hPF4 tetramers and crosslinking the DNA into higher molecular weight complexes, sterically preventing APS antibodies from binding to β2GPI within the complexes.

Anti-hPF4 moAbs, G4KKO, and RTO, were effective in preventing the enhanced neutrophil rolling in the passive APS murine model if given prophylactically (pre-APS IgG), but less so, if given therapeutically (post-APS IgG). In contrast, thrombus formation was improved when G4KKO or RTO was given either prophylactically or therapeutically. We propose that these observations are due to the enhanced rolling of neutrophils on the endothelium being too intense and occurring too rapidly to be interdicted once initiated. It may be necessary to modify the model to cause a more chronic state to explore the long-term consequences of blocking PF4 alone or in combination with anticoagulants to address the potential to disrupt ongoing, prothrombotic processes in APS patients. Also, why platelet thrombosis was more responsive to therapeutic intervention than neutrophil rolling and neutrophil incorporation into thrombi is unclear. Platelet thrombosis, and neutrophil and endothelial activation involve overlapping, but likely also distinct pathways and may also differ in the role of FcγRIIA and complement activation.

Our data also suggest that VWF may be involved as well as NETs in the prothrombotic state of APS, just as we showed for HIT^36^. Whether there are PF4:β2GPI:VWF immune complexes as well as PF4:β2GPI:NET immune complexes in APS, or whether our findings in **Figure *S*7** are due to interactions of NETs with VWF strands so that both are necessary for optimal thrombus stability will need further study.

The murine studies provide insights into the “joint biology” of PF4 and β2GPI. Unlike in mPF4^-/-^ mice, thromboinflammation was seen in WT mice injected with human APS IgG as well as in the hPF4^+^ and HIT mice, indicating that hPF4, as well as mPF4, likely complexes mβ2GPI on NETs to be recognized by patient anti-hβ2GPI antibodies. Mature hPF4 and mPF4 only share approximately 60% homology^43^, suggesting that both species of PF4 bind to mβ2GPI through their better-conserved C-termini and interact with a similarly conserved region between hβ2GPI and mβ2GPI, perhaps in their Domain III as previously proposed^11,44,45^.

The biological pressure driving parallel conservation for PF4 and β2GPI is unclear. Both PF4 and β2GPI have a long list of possible biological functions^9,46^, but one common and important biological function is that both proteins are needed to protect mice against microbial sepsis^29,47–49^. We have shown that PF4:NETs are important in protecting against sepsis by clearing circulating microbes and endotoxic products^29^. We have proposed that protection against microbial sepsis is the key force driving the presence of PF4, a highly expressed chemokine released by activated platelets with uniquely high affinity for various polyanions, ideal for removing large numbers of microbes and microbial degradation products onto NETs^29^. We propose that β2GPI contributes to this protective effect and that PF4:β2GPI:NET complexes are involved in this pathway to protect against microbial-induced diffuse multi-organ injury. Additional studies will be needed to confirm if the two proteins complexed to NETs cooperate in protection against sepsis.

The extent of thrombosis after infusion of APS IgG in HIT mice was similar to that seen in littermate hPF4^+^ mice with the exception of neutrophil and neutrophil EV incorporation. Incorporation of intact neutrophils, which was more extensive in the venule thrombi of the FcγRIIA-expressing mice as was incorporation of neutrophil-derived EVs in arteriole thrombi, although both did not reach statistical difference. Prior studies had not definitively shown a clear role for FcγRIIA activation in APS thromboinflammation^50^. Our data suggest that activation through FcγRIIA may modestly enhance neutrophil activation in both venule and arteriole thrombi with even less of an impact on platelet and fibrin incorporation. The mechanistic basis for this distinction and its clinical impact of blocking the Fc pathway in APS remains unclear.

Studies of additional triple-positive APS patients with a history of thrombosis beyond the four reported here are needed to affirm our conclusion that PF4 contributes to immunothrombosis. Additional studies are also needed to compare plasma levels of PF4 and, if possible, circulating products from PF4:β2GPI:NETs, in individuals with IgG antibodies and thrombosis *vs.* patients without a history of thrombosis to see if these levels predict thrombotic or adverse gestational complications. In murine models of APS in pregnancy, β2GPI is concentrated in the placenta in the maternal vasculature and at sites of implantation^51^, but the presence of PF4:β2GPI:NET complexes and potential relationships between placental accumulation and complications of pregnancy has not been examined. Whether placental pathology and pregnancy loss will develop in APS IgG-immunized mPF4^-/-^ mice is unclear, as is whether gestational complications are greater in HIT than in hPF4^+^ mice lacking FcγRIIA.

In summary, we examined the role of PF4:β2GPI:NET complexes in APS. Our data show the formation of large PF4:β2GPI:NET antigenic complexes that bind anti-β2GPI APS antibodies to form immune complexes that promote thromboinflammation both *in vitro* in microfluidic systems and *in vivo* in mice. hPF4 or mPF4, is needed in a passive infusion model of APS IgG to develop thromboinflammation. Each of three mechanistically distinct, Fc-modified anti-hPF4 moAbs blocked APS IgG-induced thromboinflammatory findings *in vitro.* The therapeutic efficacy of blocking PF4 biology in APS were confirmed for the two anti-hPF4 moAbs tested when given as a single agent both prophylactically and therapeutically in mice post APS-IgG infusion. Our studies support a role for PF4:β2GPI:NET complexes in APS immunothrombosis and offer a novel mechanism-based therapeutic strategy that may lessen the intensity and/or duration of need for anticoagulants in APS management.

## Supporting information

Neutrophil rolling pre-APS1 IgG infusion into a WT mouse

Neutrophil rolling post-APS1 IgG infusion into a WT mouse

Neutrophil rolling pre-APS1 IgG infusion into an hPF4+mPF4-- mouse

Neutrophil rolling post-APS1 IgG infusion into an hPF4+mPF4-- mouse

Neutrophil rolling pre-APS1 IgG infusion into mPF4-- mouse

Neutrophil rolling post-APS1 IgG infusion into mPF4-- mouse

Cremaster venule thrombosis in a HIT mouse post-APS1 infusion

Cremaster venule thrombosis in a mPF4-- mouse post-APS1 infusion

Cremaster arteriole thrombosis in a HIT mouse post-APS1 infusion

Cremaster arteriole thrombosis in a mPF4-- mouse post-APS1 infusion

## Acknowledgments

This work was supported by grants from the National Heart, Lung, and Blood Institute to MP, LR and DBC (R35HL150698), to GMA, DBC, LR and MP (R01HL151730) and to KG (R00HL177830). CF was supported by an American Society of Hematology Graduate Award (GRT-40089313). AS was supported by a Hemostasis and Thrombosis Research Society grant (GRT-00005101). We thank Dr. Leslie Raffini at the Children’s Hospital of Philadelphia for providing the impetus to begin to pursue these studies.

## Contribution to this manuscript

COF and AS share first authorship and carried out the majority of the murine studies and microfluidic studies, respectively. KB carried out and interpreted the dynamic light scattering studies. These three also participated in drafting the initial and subsequent manuscript iterations. HK, SKY, JO, GZ, SJ and MG contributed to the development and performance of assays for the *in vitro* studies and/or assisted with the *in vivo* studies and/or the creation and expression of G4KKO. KRM and TLO provided antiphospholipid patient samples, insights into the nature of this disease and valuable feedback on the manuscript. GMA provided the monoclonal antibodies KKO, TRA and RTO and had helpful discussions about this manuscript. JR and YG provided 1E12. MAK carried out initial murine studies and had helpful discussions about this manuscript. DBC participated in discussions related to the direction and interpretation of experimental outcomes, and in editing the manuscript. KG provided important guidance into the neutrophil-lined microfluidic studies and carried out early studies with that system. She also participated in discussions related to this manuscript and prepared the artwork. LR characterized G4KKO biology. She and MP led the overall study design, data analysis, and manuscript preparation. MP developed the underlying concepts in this manuscript.

