## Supplemental Methods and Figures for "Antiphospholipid syndrome (APS) is a platelet factor 4 (PF4)-centric immunothrombotic disorder"

### Supplemental materials and methods

*Construction and characterization of G4KKO*

G4KKO is a chimeric monoclonal antibody engineered from the murine anti-PF4/heparin antibody KKO[1]. Using the published amino acid sequence of the KKO Fab fragment (Protein Data Bank under accession code 4R97), the DNA sequences of the most homologous variable region genes – AF025443 and EU159568 for the heavy chain variable region (VH), and AY005823 for the light chain variable region (VL) – were selected as templates. These codon sequences were modified to match the amino acid sequence of KKO. To minimize Fcγ receptor-mediated cell activation, the murine IgG2b constant regions were replaced with the human IgG4 constant region (GenBank: AF237586). The light chain was constructed by fusing the murine KKO VL sequence with the human kappa constant region, using the sequence from NCBI EF589452, referenced against NCBI AB289328. (**Figure *S*1A**). Both hybrid heavy and light chain genes were synthesized (Integrated DNA Technologies) and cloned into the pMT vector (ThermoFisher) for expression in S2 insect cells, with a BiP signal sequence to promote secretion into the medium[2]. After confirming secretion of PF4/heparin-binding human IgG, the antibody was purified by Protein G affinity chromatography (ThermoFisher)[3], and its purity verified by SDS-PAGE.

Binding specificity was assessed by ELISA. Plates were coated with PF4/heparin complexes (10 µg/ml PF4 + 0.2 U/ml heparin), and specificity was confirmed by inhibition with excess heparin (100 U/ml) in control wells. Binding was detected using horse-radish peroxidase-conjugated anti-human IgG antibody (Jackson ImmunoResearch) (**Figure *S*1B**). The nonlinear fit of specific binding was determined using the Hill equation (Prism 10.6.0).

Platelet and complement activation by G4KKO were measured in whole blood by flow cytometry. Diluted blood (human or mouse from transgenic mice expressing FcγRIIA[4]; 1:50 v/v) was incubated with indicated amount of KKO or G4KKO (0-200 µg/ml) in the presence of PF4 (10 µg/ml) for 45 min. Platelet activation was measured by expression of P selectin (anti-hCD62P, BD Biosciences; anti-mCD62P, Emfret Analytics) and complement activation by deposition of complement on human platelets (anti-hC3c, Abcam) (**Figure *S*1C- *S*1E**).

**Supplemental videos**

**Video #1. Neutrophil rolling pre-APS1 IgG infusion into a WT mouse.**

**Video #2. Neutrophil rolling post-APS1 IgG infusion into a WT mouse.**

**Video #3. Neutrophil rolling pre-APS1 IgG infusion into an $hPF4^{+}/mPF4^{-/-}$ mouse.**

**Video #4. Neutrophil rolling post-APS1 IgG infusion into an $hPF4^{+}/mPF4^{-/-}$ mouse.**

**Video #5. Neutrophil rolling pre-APS1 IgG infusion into $mPF4^{-/-}$ mouse.**

**Video #6. Neutrophil rolling post-APS1 IgG infusion into a $mPF4^{-/-}$ mouse.**

**Video #7. Cremaster venule thrombosis in a HIT mouse post-APS1 infusion.**

**Video #8. Cremaster venule thrombosis in a $mPF4^{-/-}$ mouse post-APS1 infusion.**

**Video #9. Cremaster arteriole thrombosis in a HIT mouse post-APS1 infusion.**

**Video #10. Cremaster arteriole thrombosis in a $mPF4^{-/-}$ mouse post-APS1 infusion.**

### Supplemental figure legends

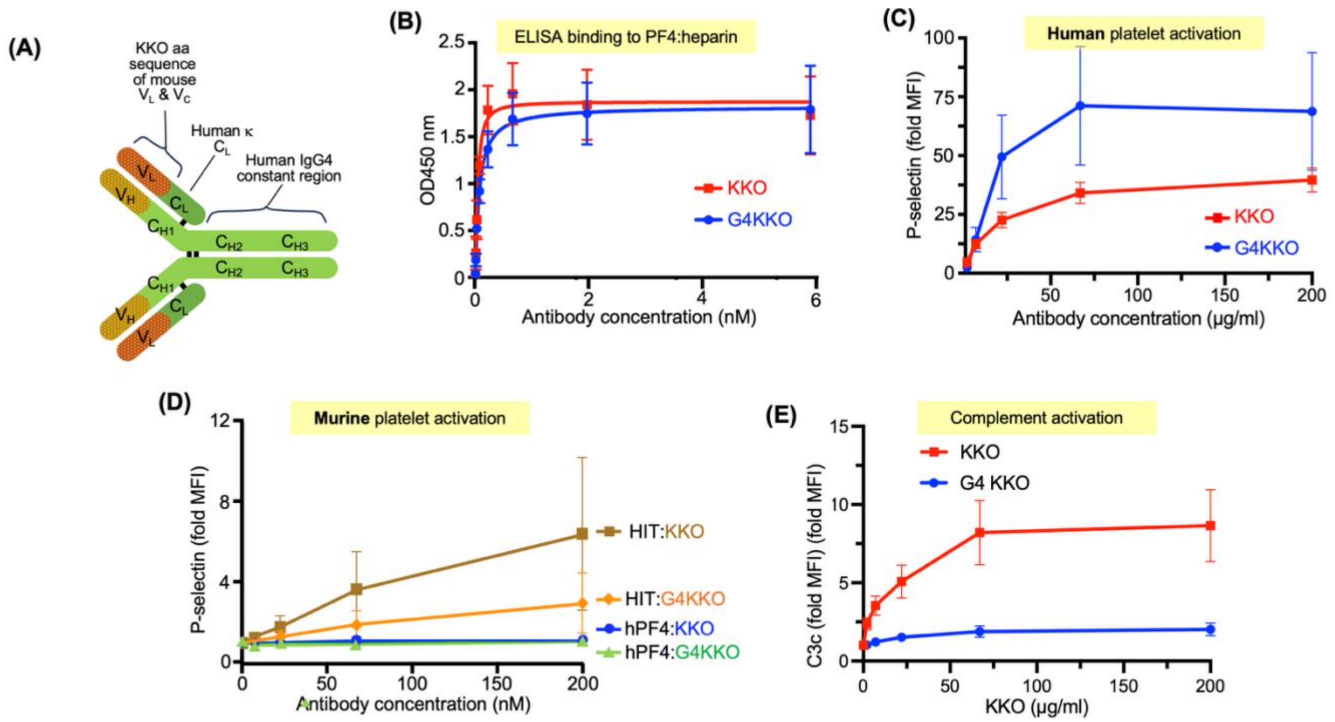


**Figure *S*1. Construction and characterization of G4KKO.** (**A**) Schematic construct of G4KKO showing the source of the various domains of G4KKO. (**B**) Graph showing best fit ± standard deviation (SD) for N = 3 experiments per arm, done in duplicate, for binding of KKO and G4KKO to hPF4:heparin-coated wells. $Kd_{KKO}$ = 0.05 nM (confidence interval 0.03-0.07); $Kd_{G4KKO}$ = 0.08 nM (confidence interval 0.05-0.14). (**C**) Human platelet activation by KKO, and G4KKO as indicated by P-selectin surface expression. Shown is mean ± SD for N=3 experiments, done in duplicate. (**D**) Same as (C), but for activation of mouse platelets. Mice were either $hPF4^{+}$ mice (lacking FcγRIIA) or HIT mice (expressing FcγRIIA). (**E**) Similar studies as in (C), but for complement C3c generation and deposition on human platelets.

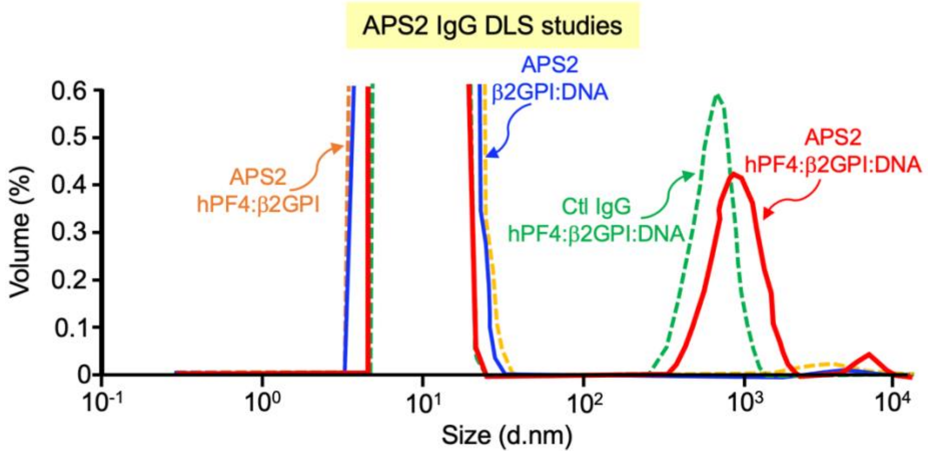


**Figure *S*2. DLS studies in the presence of β2GPI, hPF4, DNA and APS antibodies.**

Similar studies as in **Figure 1B**, but using APS2 and complexes of β2GPI:DNA, hPF4:β2GPI, and hPF4:β2GPI:DNA.

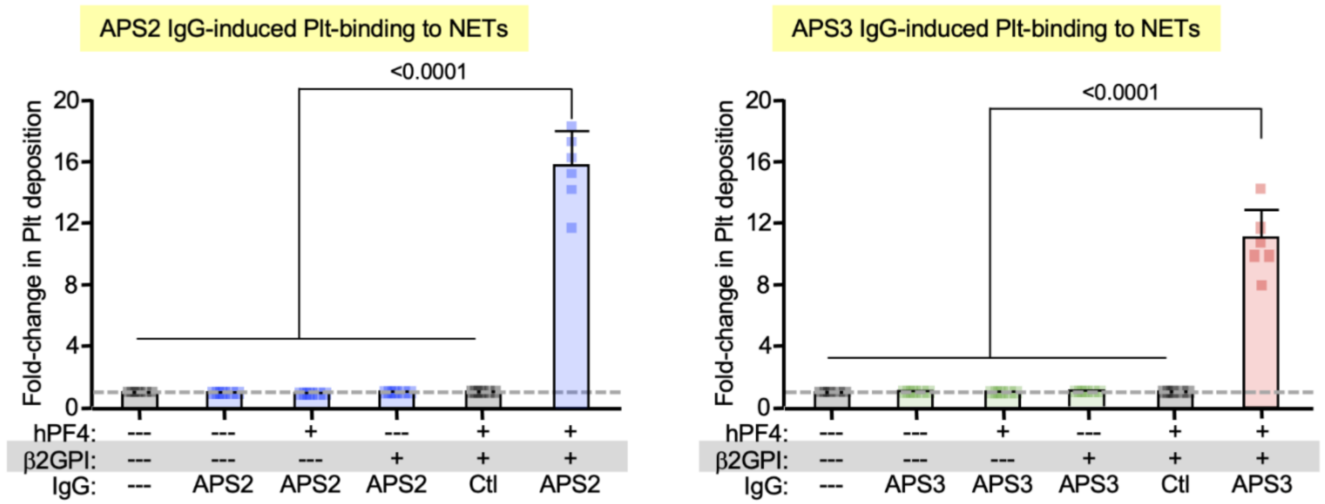


**Figure *S*3. APS IgGs enhance platelet binding to NETs only when hPF4 and β2GPI are both present.**

Similar to **Figure 3B**, but showing studies with APS2 (left) and APS3 IgGs (right).

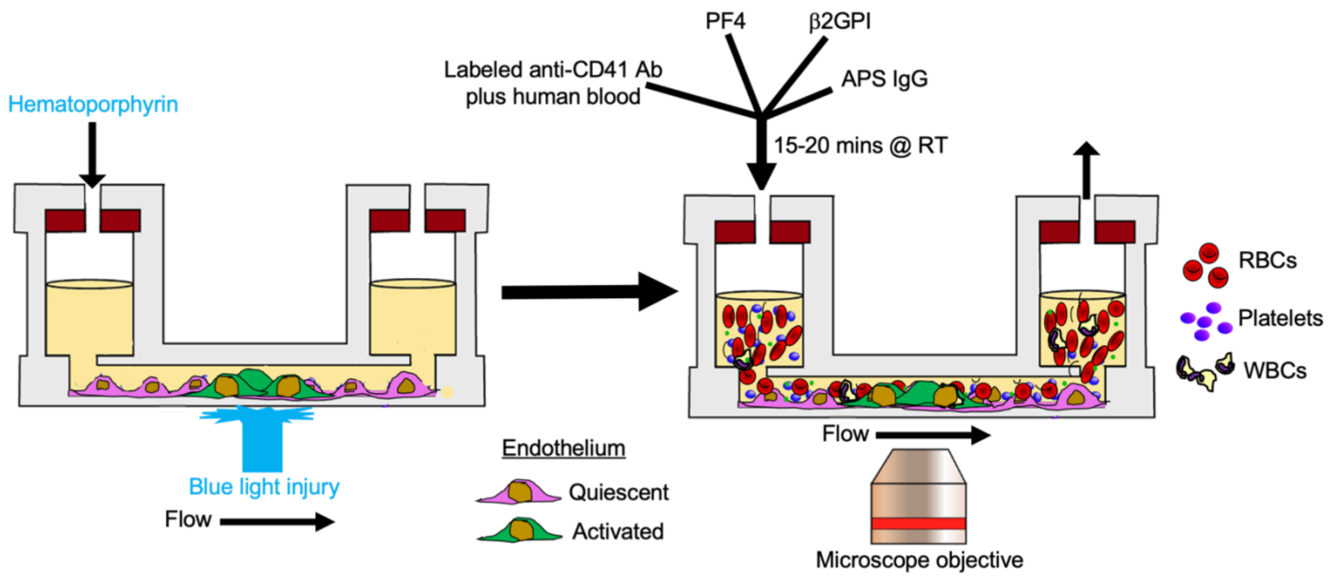


**Figure *S*4. Schematic of the photochemical injury platelet-accumulation studies of HUVEC-lined microfluidic channels in APS.**

Design of the study to use injured HUVEC-lined microfluidic channels to study the importance of hPF4 and β2GPI for thrombosis in blood exposed to IgGs isolated from patients with triple-positive APS. On the left is the HUVEC-lined channel exposed to light while hematoporphyrin is infused through the channel, injuring the HIVECs. On the right is the subsequent perfusion of whole blood supplemented with labeled, anti-CD41 antibody ± hPF4 ± β2GPI ± APS or control IgG over 15 minutes, recording the deposition of platelets, fibrin and complement.

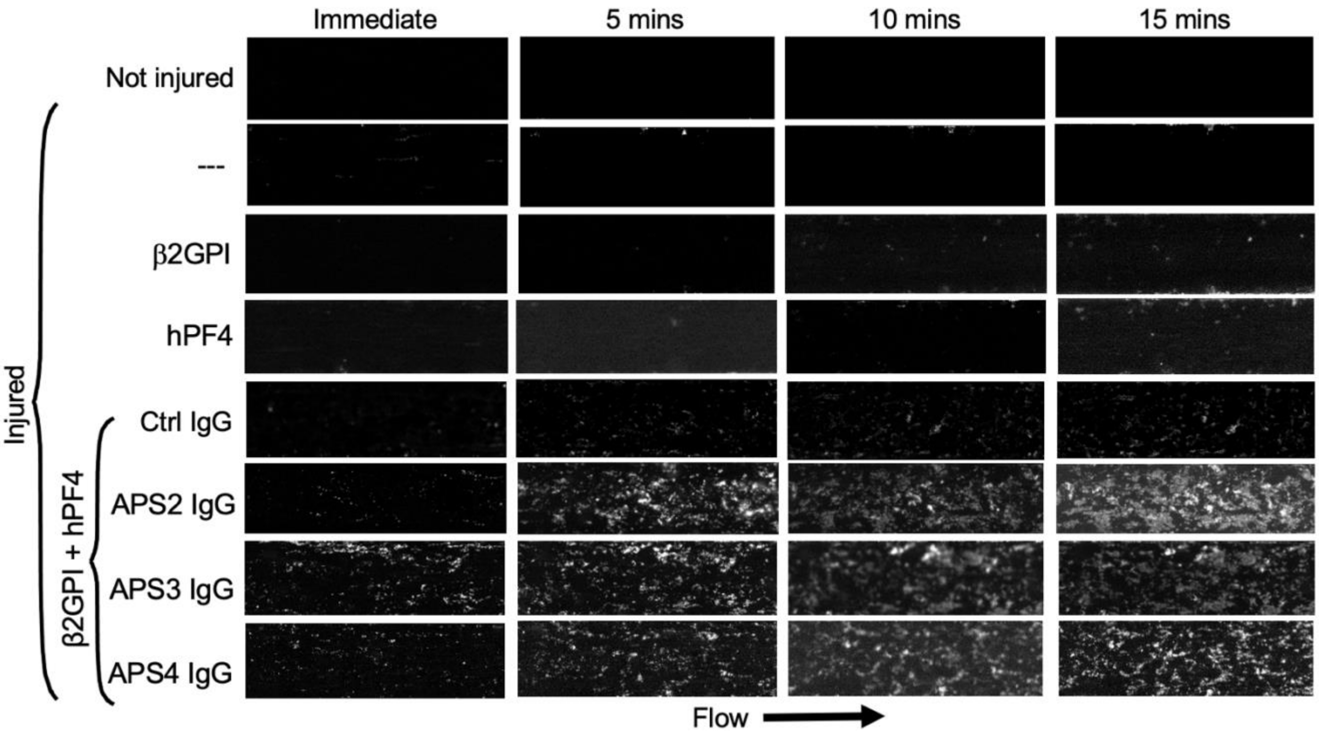


**Figure *S*5. Supplemental photochemical injury platelet-accumulation studies of HUVEC-lined microfluidic channels in APS.**

Similar to **Figure 4A** in the manuscript using APS1 IgG, but showing studies with APS2 through APS4 IgGs.

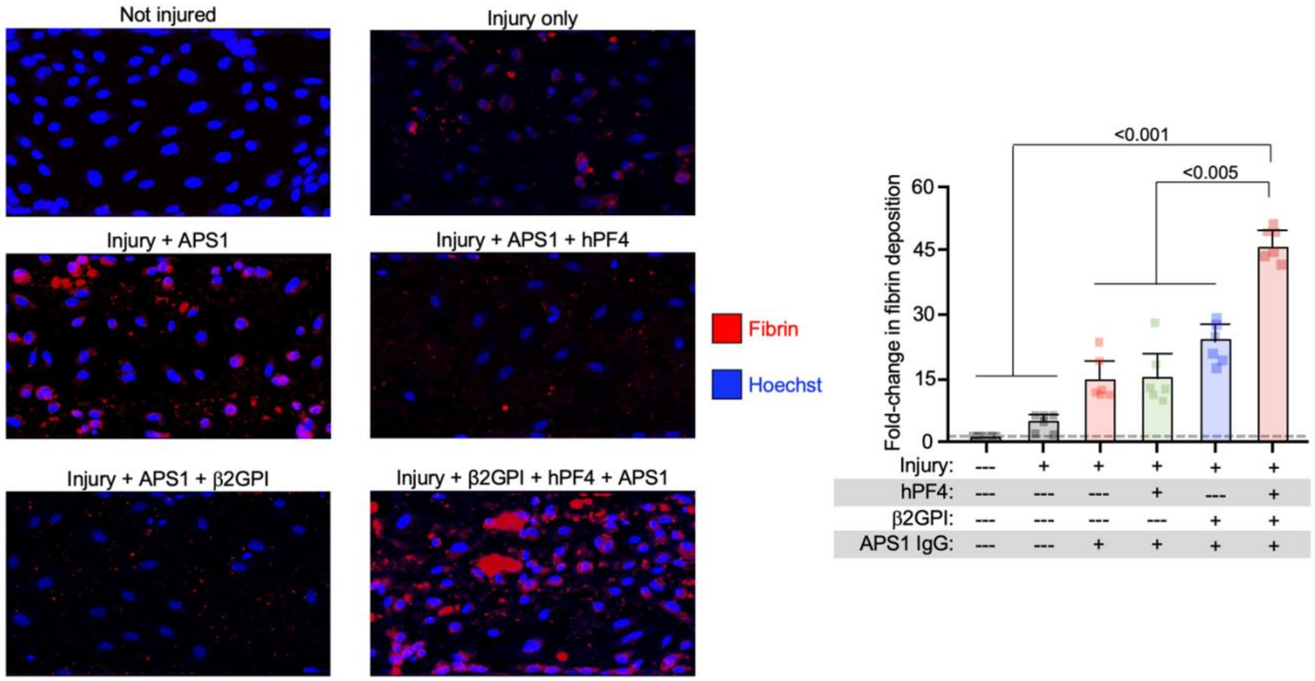


**Figure *S*6. Fibrin deposition in the photochemical-injured HUVEC thrombosis model in APS.**

Left: Representative images of channels lined with injured HUVECs with the endothelial cell nuclei stained blue with Hoechst in studies similar to those in **Figure 4A**. Fibrin was stained with Alexa Fluor561-labeled anti-fibrin antibody. Right: Mean ± 1 SD of the fold-change in fibrin stain compared to fibrin deposition on uninjured HUVECs. N = 6 separate studies each done in duplicate. Statistical analysis was done using two-way ANOVA compared to the uninjured studies.

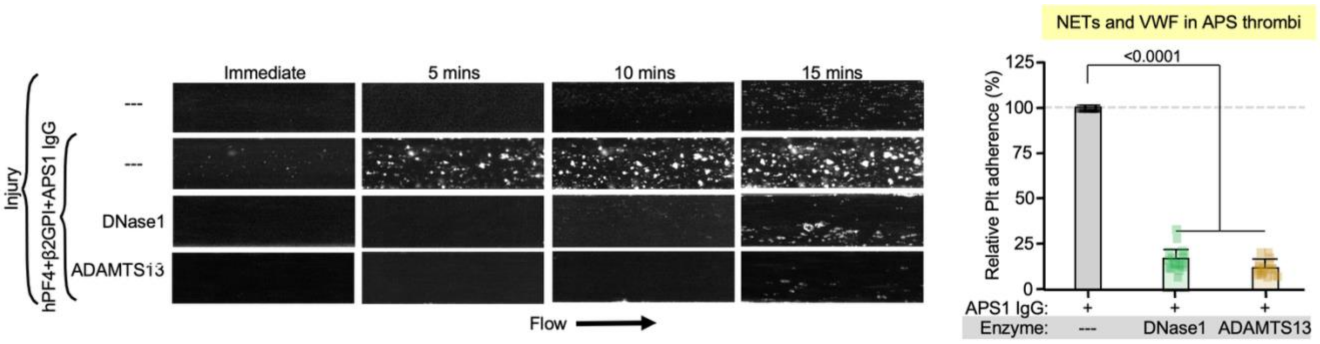


**Figure *S*7. Photochemical- injured HUVEC microfluidic studies in the absence or presence of ADAMTS13 or DNase1 in APS1 IgG-induced NET release.**

<u>Left</u>: Representative study as in **Figure 4A**, but with either DNase1 or ADAMTS13 added. <u>Right</u>: Mean ± 1 SD comparing platelet adherence to injured HUVEC channels with or without added DNase 1 or ADAMTS13 showing relative remaining platelets adherent to the vessel after treatment. N = 3 studies done in each of 3 health control’s blood in duplicate. Grey dashed line is value seen without added DNase 1 or ADAMTS13 and set to 100%. P values were determined by two-way ANOVA compared to the absence of both DNase1 or ADAMTS13.

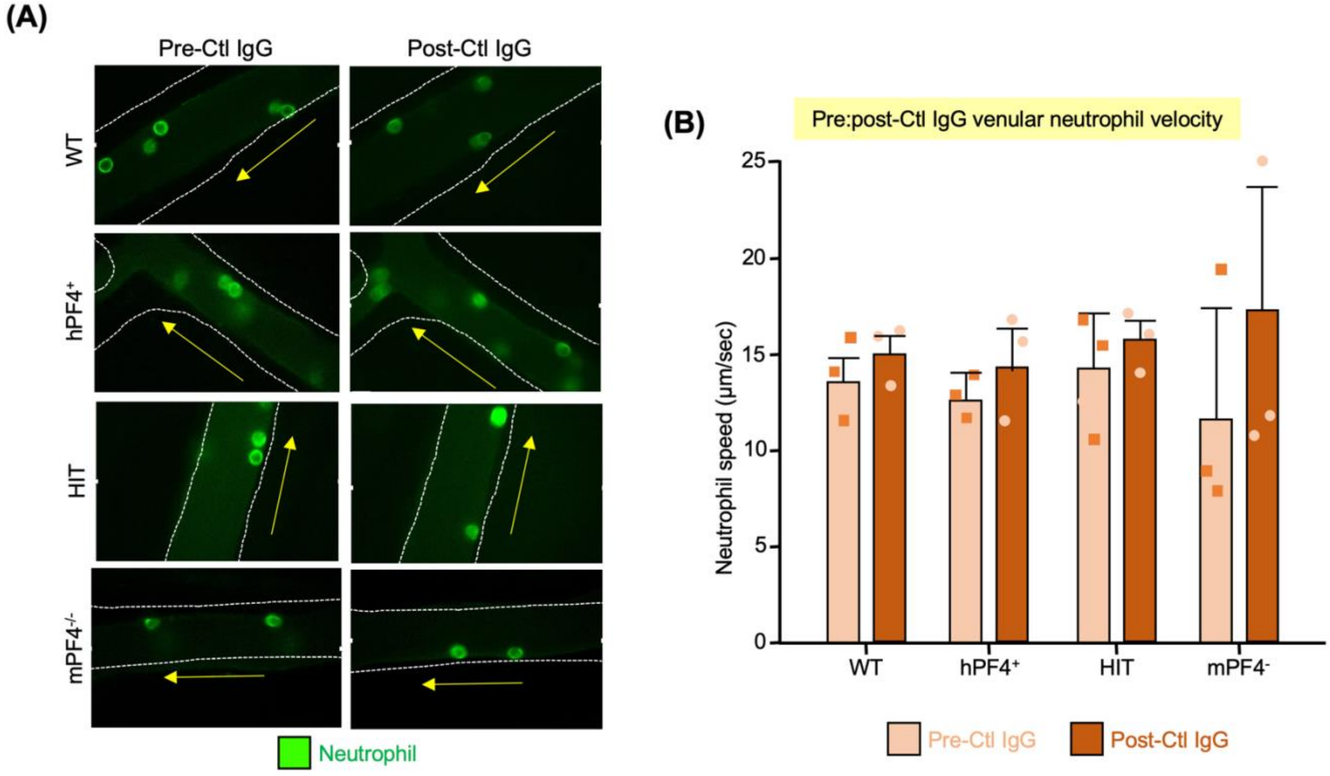


**Figure *S*8. Neutrophil rolling on cremaster venules after healthy control IgG infusions.**
Similar studies as in **Figure 5**, but with control (Ctl) IgG from a healthy individual rather than APS1 IgG. (**A**) Representative images of neutrophil cremaster venular rolling looking at the identical vein pre- and post-control IgG IV infusion. The direction of venous flow is indicated by a yellow arrow and the vessel walls outlined by a white dashed line. (**B**) Mean ± 1 SD for neutrophil rolling speed pre- and post-IgG infusion. N = 3 studies per arm. P values were determined by two-way Student t-test in each genotype mouse of pre-to-post values.

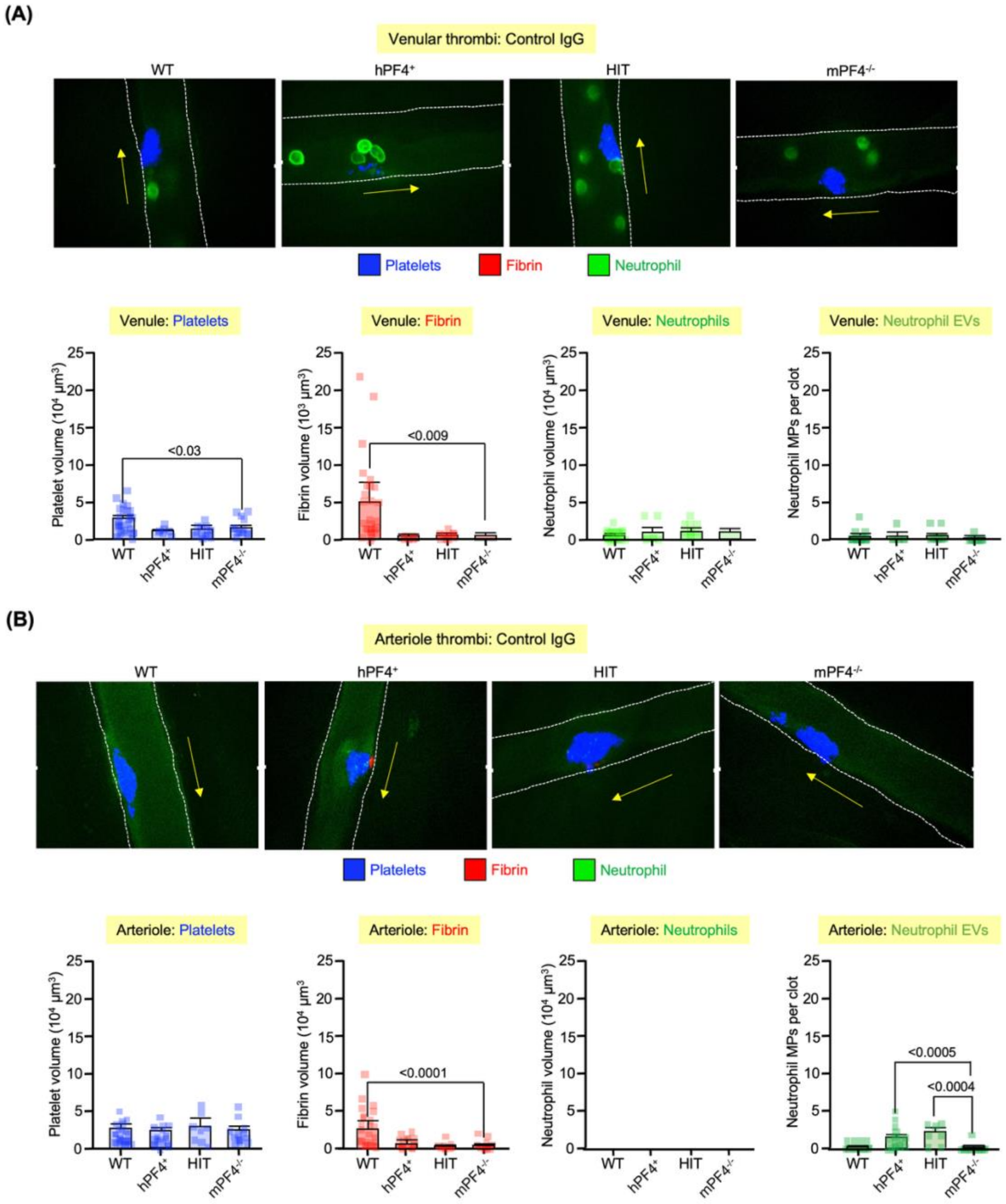


**Figure *S*9. Thrombus formation in a passive-infusion murine model infusing control IgG from healthy individuals.**

Similar to the studies in **Figure 6**, but with human control rather than APS IgG. (**A**) are venous studies and (**B**) arteriole studies. Mean ± 1 SD with 1-4 mice were studied per arm with 8-32 injury per studied mouse. Statistics done by two-way ANOVA.

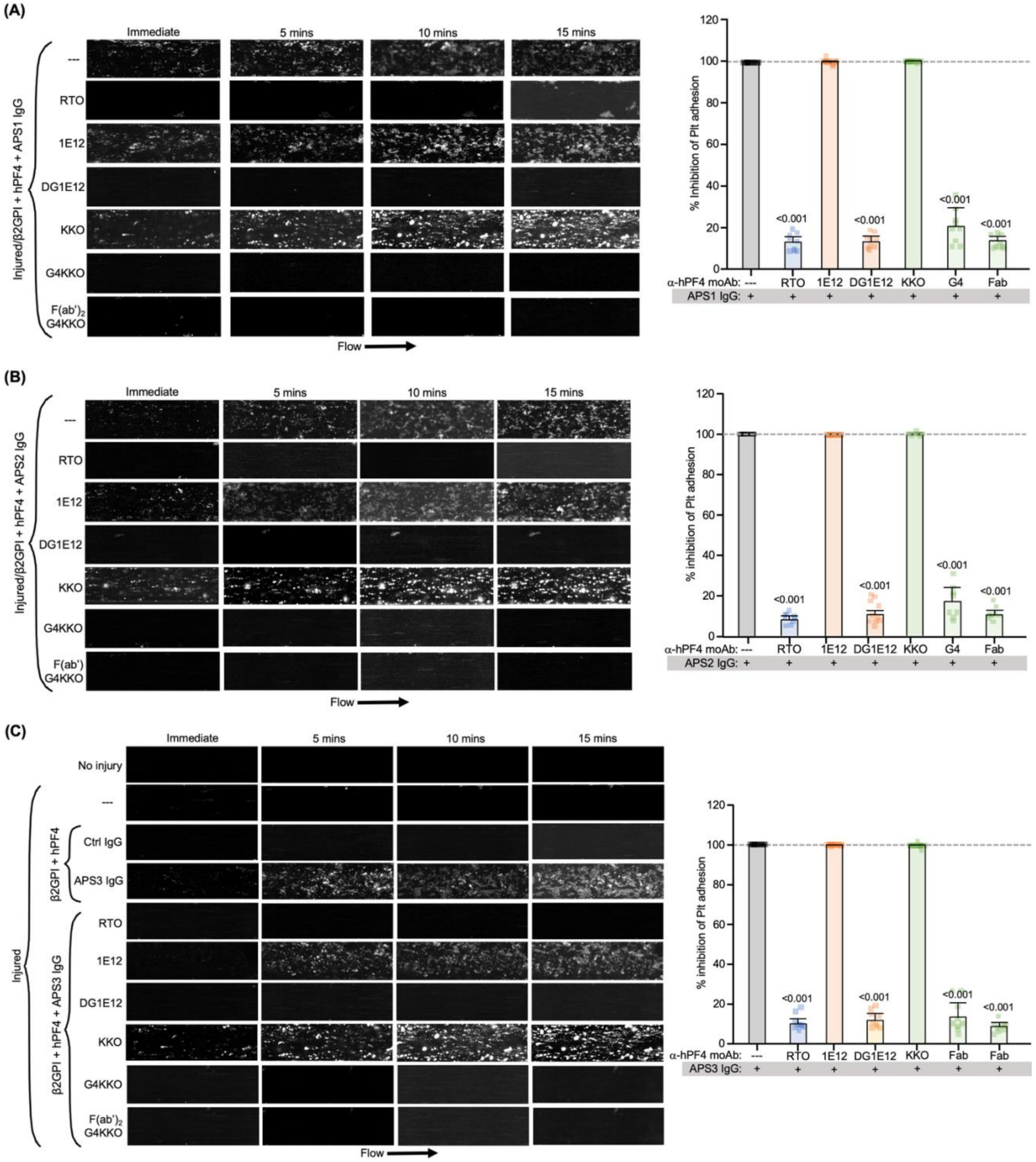


**Figure *S*10. The effects of the anti-hPF4 moAbs on APS IgG-induced thrombosis in the injured HUVEC microfluidic system.**

Similar studies as in **Figure 4**, but with added anti-hPF4 moAbs as indicated. (**A**) thru (**C**) are studies with three different triple-positive anti-β2GPI IgG preparations, APS 1 thru APS3, respectively. Left: Representative images in the absence and presence of the indicated anti-hPF4 moAbs. In (C), two additional controls are shown indicating that few platelets adhered if the HUVECs are not injured and that inclusion of IgG isolated from a healthy control also does not lead to significant platelet adherence. Right: Mean ± 1 SD for each anti-PF4 blocking moAb. N = 4, each done in duplicate, for each APS IgG studied. Dashed line is the mean level of platelet thrombosis seen with the APS IgG alone. Analyzed by two-way ANOVA. Significant differences from no added blocking antibody bar at the left in grey are shown.

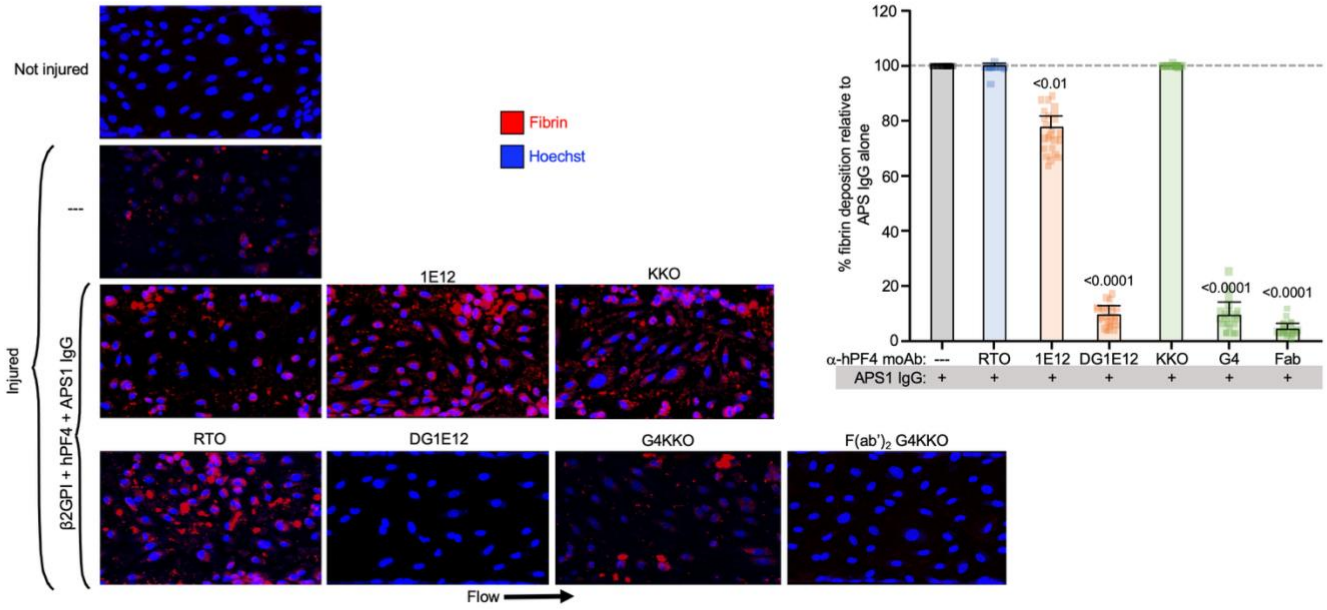


**Figure *S*11. Ant-hPF4 moAbs effects on fibrin incorporation in the photochemical-injured HUVEC microfluidic model of APS.**

Left: Representative images of fibrin incorporation into injured HUVEC microfluidic channels as in **Figure 4** using APS1 IgG. Images shown were done after the 15-minute endpoint of the platelet thrombosis studies. Right: Mean ± 1 SD of fibrin accumulation as in **Figure *S*5**. N = 4 for APS1 through APS3 IgGs using 3 different healthy blood donors. Individual data points are shown. P values are only shown were significant differences were seen after two-way ANOVA analysis when compared to APS IgG-alone treatment arm.

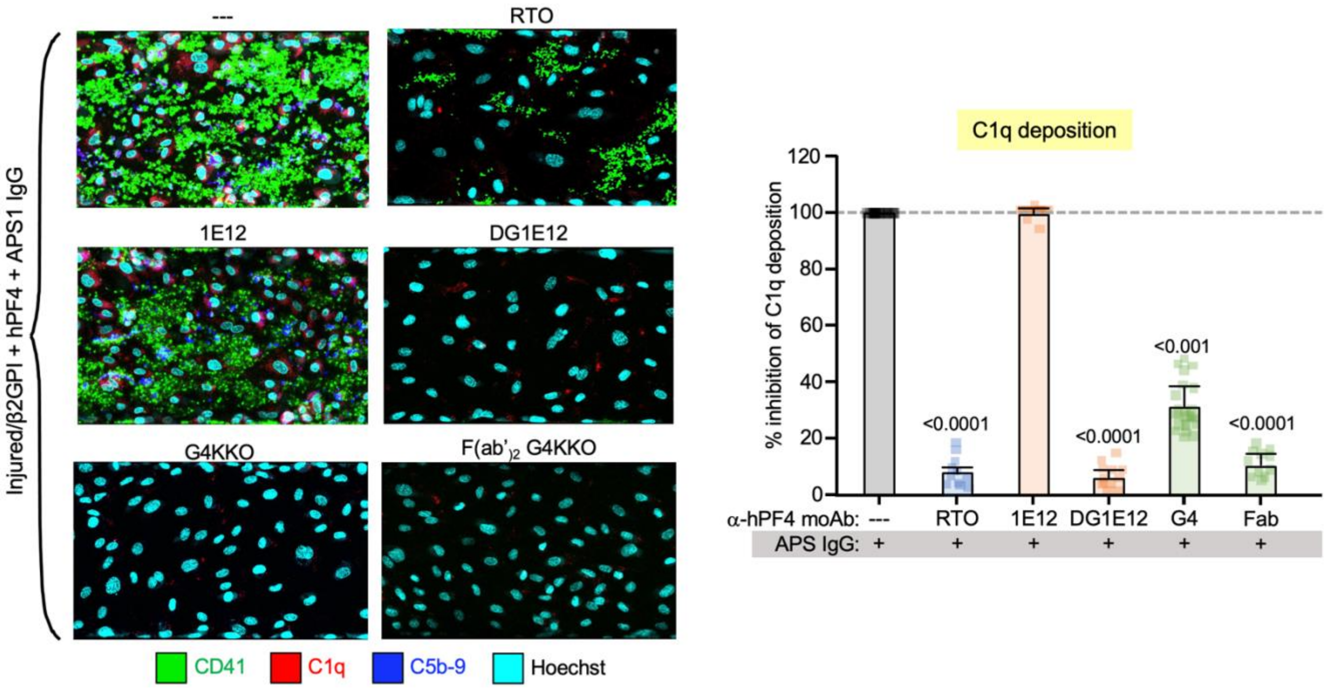


**Figure *S*12. Complement deposition in the photochemical-injured HUVEC microfluidic system after hPF4 plus β2GPI plus APS1 IgG exposure for each anti-hPF4 moAb intervention.**

Complement studies similar to **Figure 7C**. Left: Representative images showing both C1q and C5b-9 deposition on HUVECs at the end of studies similar to **Figure 4B** with platelets stained for CD41 (green), C1q (red), C5b-9 (dark blue) and HUVECs by Hoechst staining of endothelial cell nuclei (aqua blue). Right: Similar studies to **Figure 7C**, but for C1q. N = 3 APS1 IgGs studies, each done in duplicate, with 2 different healthy donor blood samples in each of the indicated anti-hPF4 moAb studies.

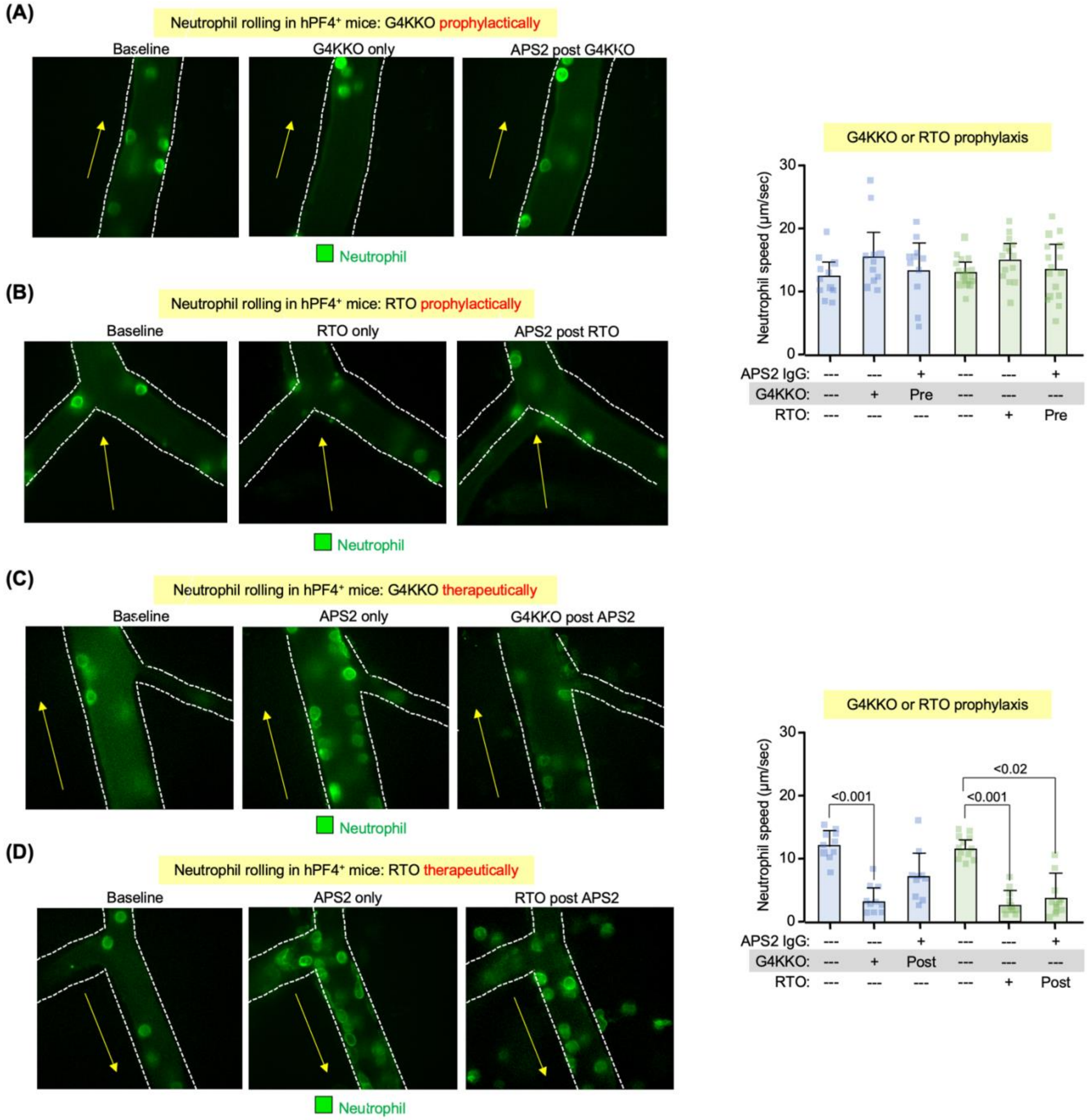


**Figure *S*13. The effects of G4KKO and RTO on neutrophil rolling: prophylactic *vs*. therapeutic intervention.**

(**A**) through (**D**) show neutrophil rolling on cremaster venules studies similar to those in **Figure 5** done in hPF4$^+$ mice receiving APS2 IgG. G4KKO or RTO were given either 15-minutes prophylactically before APS2 IgG in (A) or (C), respectively, or therapeutically 15 minutes after APS2 IgG in (B) or (D). Left: Representative images as described in **Figure 5.** Right: Mean ± 1 SD of speed of neutrophil rolling. N

= 9-24 per arm. Statistical analysis was by two-way ANOVA. P values are shown compared to the baseline arm where the mice had not received APS2 IgG or an anti-hPF4 moAb.

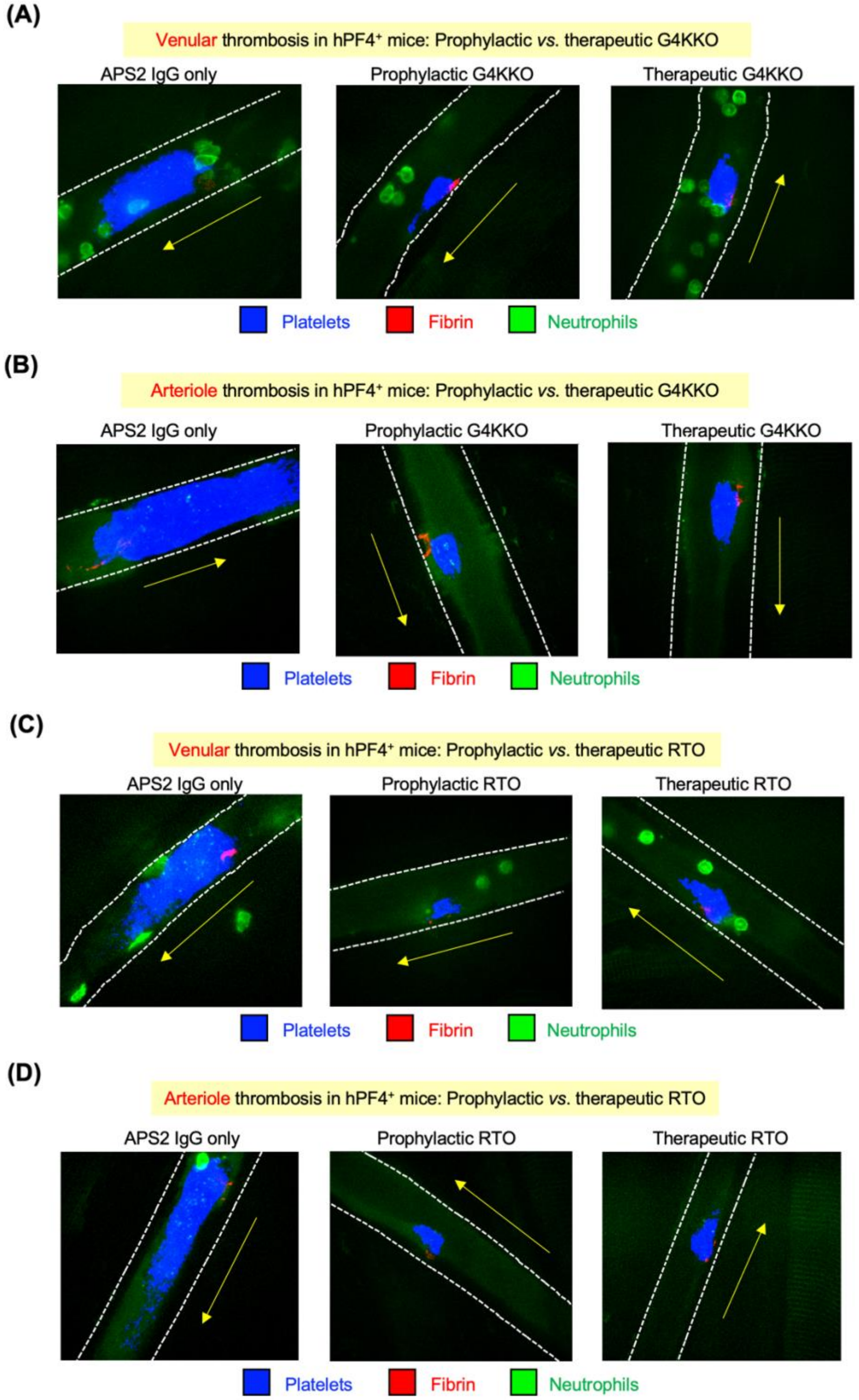


**Figure *S*14. The effects of G4KKO and RTO on thrombosis prophylactic *vs*. therapeutic intervention.**

(**A**) and (**B**) are representative images studying the use of G4KKO to inhibit thrombosis in the APS model in hPF4$^{+}$ mice. These images accompany **Figures 7D** and **7E**, respectively, and are similar to the studies presented in **Figure 6A**. G4KKO was either not given (left images) or was administered prophylactically prior to the APS2 IgG infusion (middle images) or given therapeutically after the APS2 IgG infusion (right images). (**C**) and (**D**) are similar studies to (A) and (B), respectively, but with RTO infused rather than G4KKO.

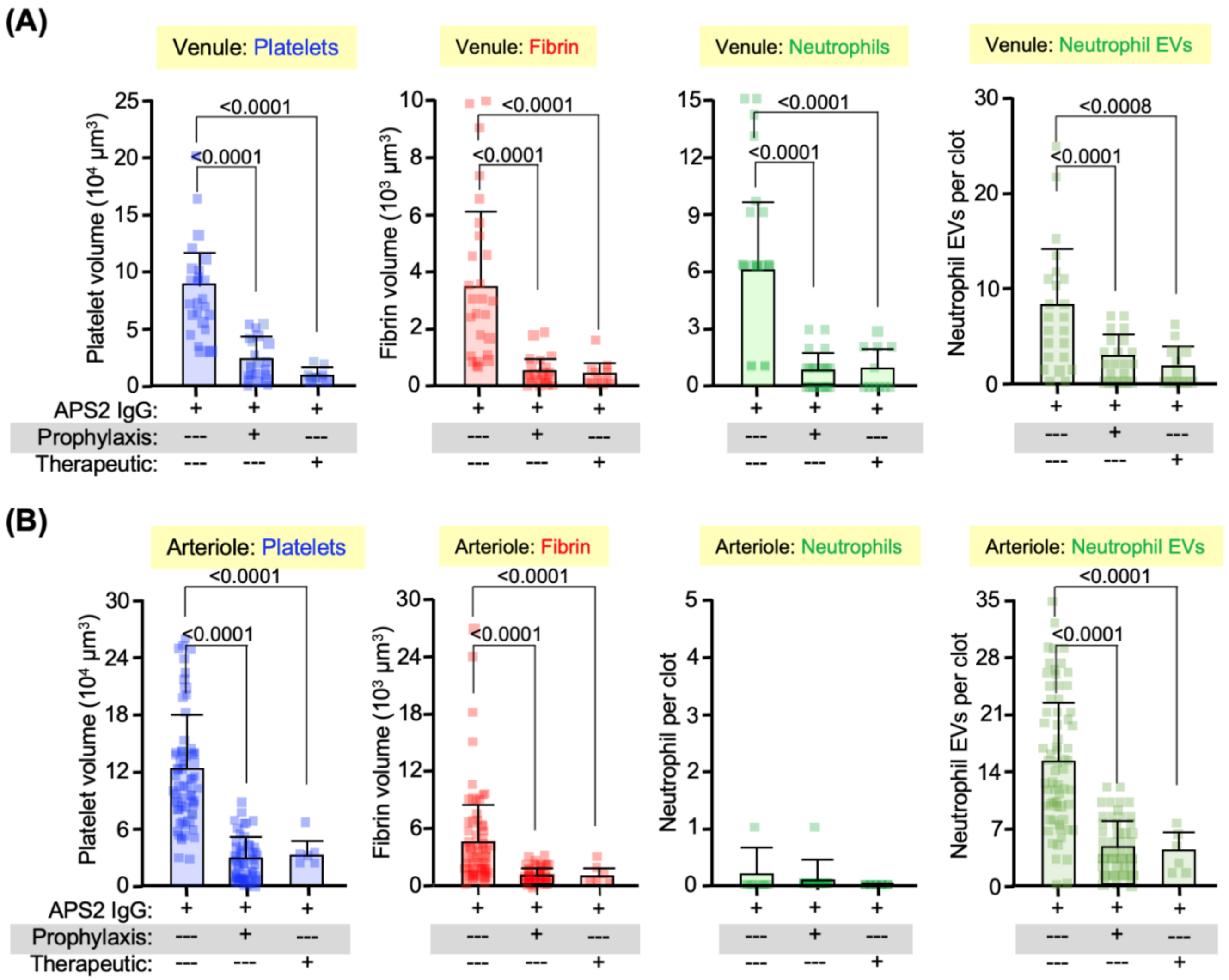


**Figure *S*15. The inhibitory effects of RTO on APS thrombosis.**

Studies were done as in **Figures 7D**, **7E** and ***S*14**, but using RTO instead of G4KKO. (**A**) are venular thrombosis and (**B**) are arteriole thrombosis. Both (A) and (B) show platelet, fibrin, neutrophil and neutrophil EV incorporation into thrombi. Mean ± 1 SD are shown as well individual study points. In (A), N = 12-26 studies mice per arm. In (B), N = 12-64 studies mice per arm. P values were determined by two-way ANOVA. Only significant comparative differences between those receiving or not receiving RTO are shown.

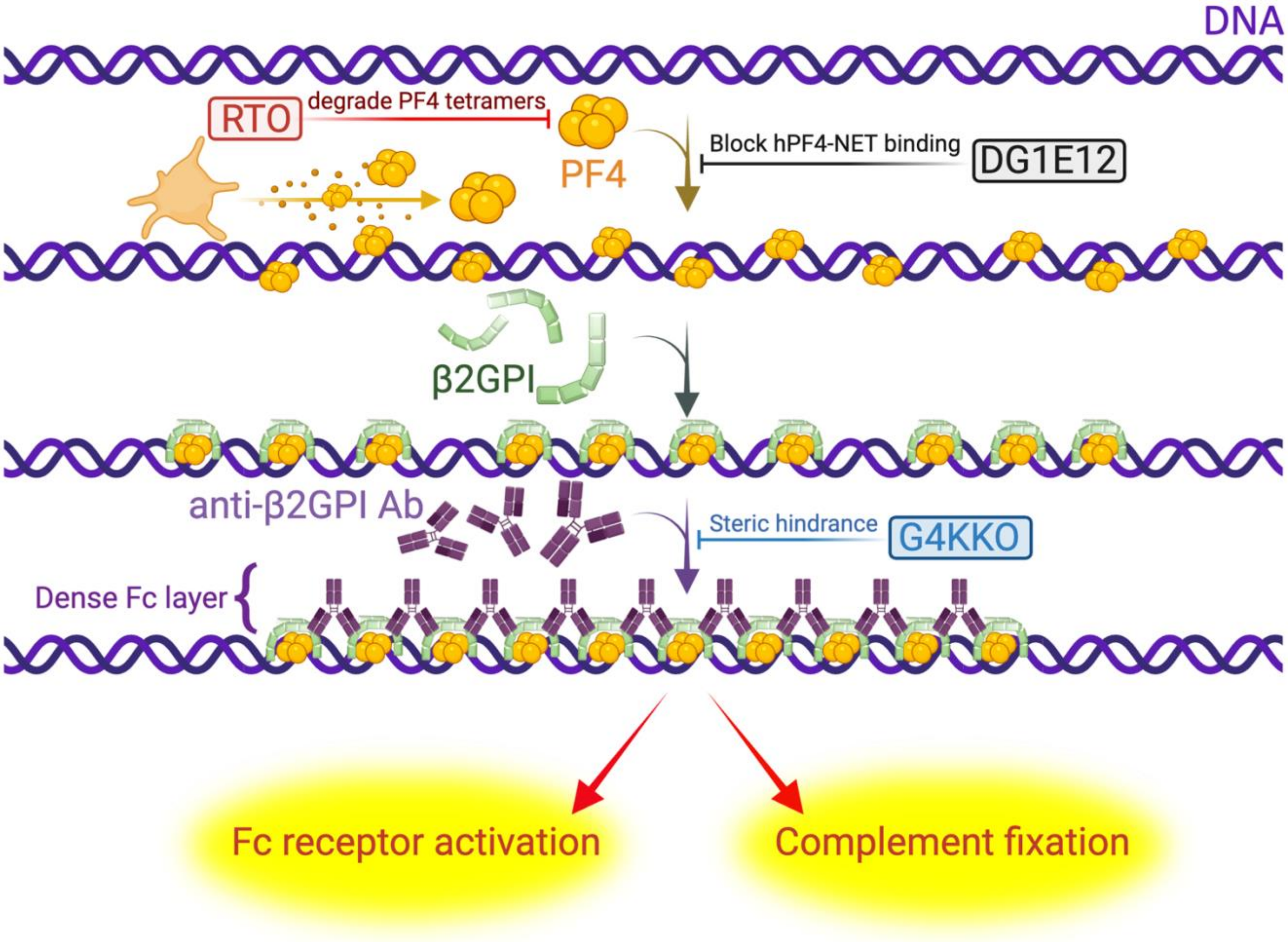


**Figure *S*16. Proposed schematic of how hPF4 and β2GPI and DNA form an APS antigenic target and the anti-hPF4 moAbs sites of inhibition.**

hPF4 tetramers bind to the phosphoribose backbone of DNA within NETs which then facilitates the binding of β2GPI, resulting in a dense antigenic target for anti-β2GPI antibodies that, in turn, activates cellular Fcγ receptors and complement. The three anti-hPF4-specific antibodies block this prothrombotic pathway at three different points: RTO blocks PF4 tetramerization so that the PF4 does not bind to the DNA. 1E12 blocks binding of hPF4 to the DNA by hiding the heparin-binding site on tetrameric PF4, and KKO sterically blocks anti-β2GPI antibodies from binding to the hPF4:β2GPI:DNA antigenic complex.
